# Structure of the Shaker Kv channel and mechanism of slow C-type inactivation

**DOI:** 10.1101/2021.09.21.461258

**Authors:** Xiao-Feng Tan, Chanhyung Bae, Robyn Stix, Ana I. Fernández-Mariño, Kate Huffer, Tsg-Hui Chang, Jiansen Jiang, José D. Faraldo-Gómez, Kenton J. Swartz

**Affiliations:** Molecular Physiology and Biophysics Section, Porter Neuroscience Research Center, National Institute of Neurological Disorders and Stroke, National Institutes of Health, Bethesda, MD 20892; Theoretical Molecular Biophysics Laboratory, National Heart, Lung and Blood Institute, National Institutes of Health, Bethesda, MD 20892; Department of Biology, Johns Hopkins University, 3400 N. Charles Street, Baltimore, MD, 21218, USA; Laboratory of Membrane Proteins and Structural Biology and Biophysics Center, National Heart, Lung, and Blood Institute, National Institutes of Health, Bethesda, MD 20892

## Abstract

Voltage-activated potassium (Kv) channels open upon membrane depolarization and proceed to spontaneously inactivate. Inactivation controls neuronal firing rates and serves as a form of short-term memory, and is implicated in various human neurological disorders. Here, we use high-resolution cryo-electron microscopy and computer simulations to determine one of the molecular mechanisms underlying this physiologically crucial process. Structures of the activated Shaker Kv channel and of its W434F mutant in lipid bilayers demonstrate that C-type inactivation entails the dilation of the ion selectivity filter, and the repositioning of neighboring residues known to be functionally critical. Microsecond-scale molecular dynamics trajectories confirm these changes inhibit rapid ion permeation through the channel. This long-sought breakthrough establishes how eukaryotic K^+^ channels self-regulate their functional state through the plasticity of their selectivity filters.

**One-Sentence Summary:** Structures of the Shaker Kv channel reveal the mechanism of slow C-type inactivation involves dilation of the selectivity filter.

---

Voltage-activated potassium (Kv) channels open and close in response to changes in membrane voltage, serving critical functions in electrical signaling in neurons and muscle, neurotransmitter and hormone secretion, cell proliferation and migration, and ion homeostasis (*1, 2*). Functional studies of Shaker Kv channels have been foundational for establishing key mechanistic principles such as voltage-sensing, pore opening and inactivation in eukaryotes (*1, 3–5*). The structure of the related Kv1.2/2.1 paddle chimera has provided a valuable framework to begin to understand these processes (*6, 7*), but it has been challenging to capture the protein in distinct conformations, thereby limiting mechanistic insights (*8, 9*). In addition, the phenotypes of well-characterized mutants in the Shaker Kv channel do not always translate to the Kv1.2 channel (*10*), leaving key questions unanswered. One of these outstanding questions is what the structural mechanism underlies the so-called C-type inactivation of Shaker Kv channels (*11*), a slow inactivation process that regulates the firing of action potentials in neurons and cardiac muscle by diminishing the availability of these channels to conduct ions. This intriguing process has been implicated in a range of neurological and psychiatric disorders (*1, 12–15*), and occurs in other Kv channels too. For example, the human ether-a-go-go related (hERG) Kv channel also enters a C-type inactivated state rapidly after opening, which translates into unique inwardly-rectifying properties that are critical for shaping the cardiac action potential and preventing arrhythmias (*16–19*). C-type inactivation has also been proposed to serve important roles in Ca^2+^-activated K^+^ channels and K2P channels (*20–22*).

It has long been hypothesized that C-type inactivation in Shaker Kv channels involves a conformational change in the external pore, as it is influenced by mutations in this region as well as by ion and blocker occupancy of the selectivity filter (*11, 14, 15, 23-35*). X-ray structures of mutants stabilizing open and inactivated states of KcsA, a prokaryotic K^+^ selective channel that consists of a pore domain homologous to that found in Kv channels, support the prevailing view that slow inactivation in K^+^ channels is caused by a collapse of the ion selectivity filter (*36–38*). However, the extent to which slow inactivation in KcsA informs on the mechanism of C-type inactivation in Kv channels remains an open question (*15, 39, 40*). In the present study we report the first-known structure of the Shaker Kv channel and that of a well-characterized mutant that dramatically accelerates C-type inactivation (*24, 30, 41, 42*). Our findings reveal that C-type inactivation in Shaker involves an unanticipated conformational rearrangement of the external pore that leads to displacement of the P-loop and dilation of the ion selectivity filter.

## Results

### Structure of the Shaker Kv channel in lipid nanodiscs

The Shaker Kv channel containing an mVenus-tag along with a deletion of residues 6-46 to remove fast inactivation (Shaker-IR) (*43, 44*) was expressed in mammalian cells, purified and reconstituted into lipid nanodiscs using MSP 1E3D1 (*8*) (**Fig. S1**), and its structure solved using cryogenic electron microscopy (cryo-EM)(**Fig. S2, S3 and Table S1**). The structural construct is identical to that used in extensive functional studies from many laboratories (*6, 11, 23-26, 28-30, 32, 35, 41, 43-47*). We determined the structure of the transmembrane (TM) regions of Shaker-IR to 3 Å resolution after focused refinement; the map was of high quality throughout, with clearly resolved densities for most side chains, making de novo atomic model building straightforward. Only two external loops within the voltage-sensing domains were unresolved (residues H254 to T276 and V330 to M356).

The fold of the tetrameric Shaker-IR Kv channel is remarkably similar to that of the Kv1.2/2.1 paddle chimera channel (*7*) (Cα RMSD 1.23Å, TM score 0.99; **Fig. S4, S5**). Each subunit in Shaker-IR is comprised of six TM helices (S1-S6), with the S1-S4 helices from each subunit forming individual voltage-sensing domains and the tetrameric arrangement of S5-S6 helices forming the central pore domain (**Figure 1A,B**). As in the Kv1.2/2.1 channel structure, Shaker-IR displays a domain-swapped architecture with the S1-S4 voltage-sensing domains positioned near the S5-S6 pore-forming helices of the adjacent subunit (**Figure 1A,B**). The cryo-EM map for Shaker-IR reveals clear densities corresponding to phospholipids that appear to be bound to the protein within both the external and internal leaflets of the lipid nanodisc (**Figure 1C,D**); many of these densities are similar to those seen in the Kv1.2/2.1 paddle chimera solved in mixed detergent/lipid micelles (*7*).

**Figure 1.**
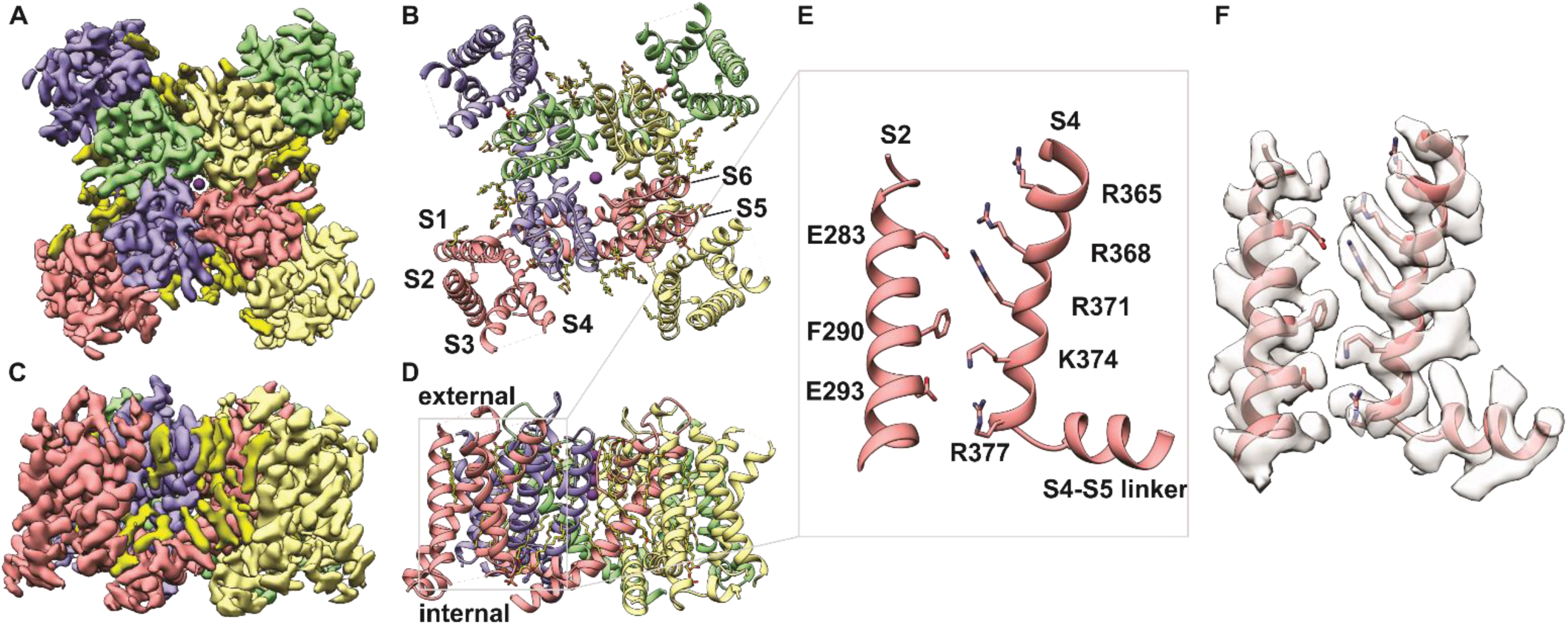
Structure of the Shaker-IR Kv channel in lipid nanodisc. **A** and **B**, View of Shaker Kv channel model (A) and EM map (B) from an extracellular perspective. **C** and **D**, Transmembrane side-view of Shaker-IR Kv channel model (C) and EM map (D). Each subunit is shown in different colors and EM densities that appear to be lipid molecules are colored in bright yellow. **E** and **F**, Close-up view of a voltage-sensing domain. The S2 and S4 helices within the voltage-sensing domain highlighted with a gray box in D is shown without (E) and with (F) EM density.

The internal structure of the S1-S4 voltage-sensing domain of Shaker-IR is also similar to that in the Kv1.2/2.1 chimera, and in both cases basic residues in S4 located near the middle of the TM region are stabilized by an occluded cation binding site or charge-transfer center that is comprised of a highly conserved Phe residue in the middle of the S2 helix (F290) and acidic residues nearby in S2 and S3 (E293 and D316) (*48*) (**Figure 1E,F**). Within the conserved core region of the S4 helix, four Arg residues (R362, R365, R368 and R371) are positioned external to F290 in S2, while one Lys (K374) is positioned near F290 and the innermost Arg (R377) and Lys (K380) are positioned internal to F290 (**Figure 1E,F**). The fact that the four basic residues in S4 known to carry most of the gating charge (*49, 50*) are accessible to the external solution suggests that our structure captures the voltage-sensing domains in the activated state, which for this channel is predominant at 0 mV. Accordingly, the intracellular S6 gate region (*51*) is open (**Figure 2C-E**), further indicating the structure represents an activated/open state.

**Figure 2.**
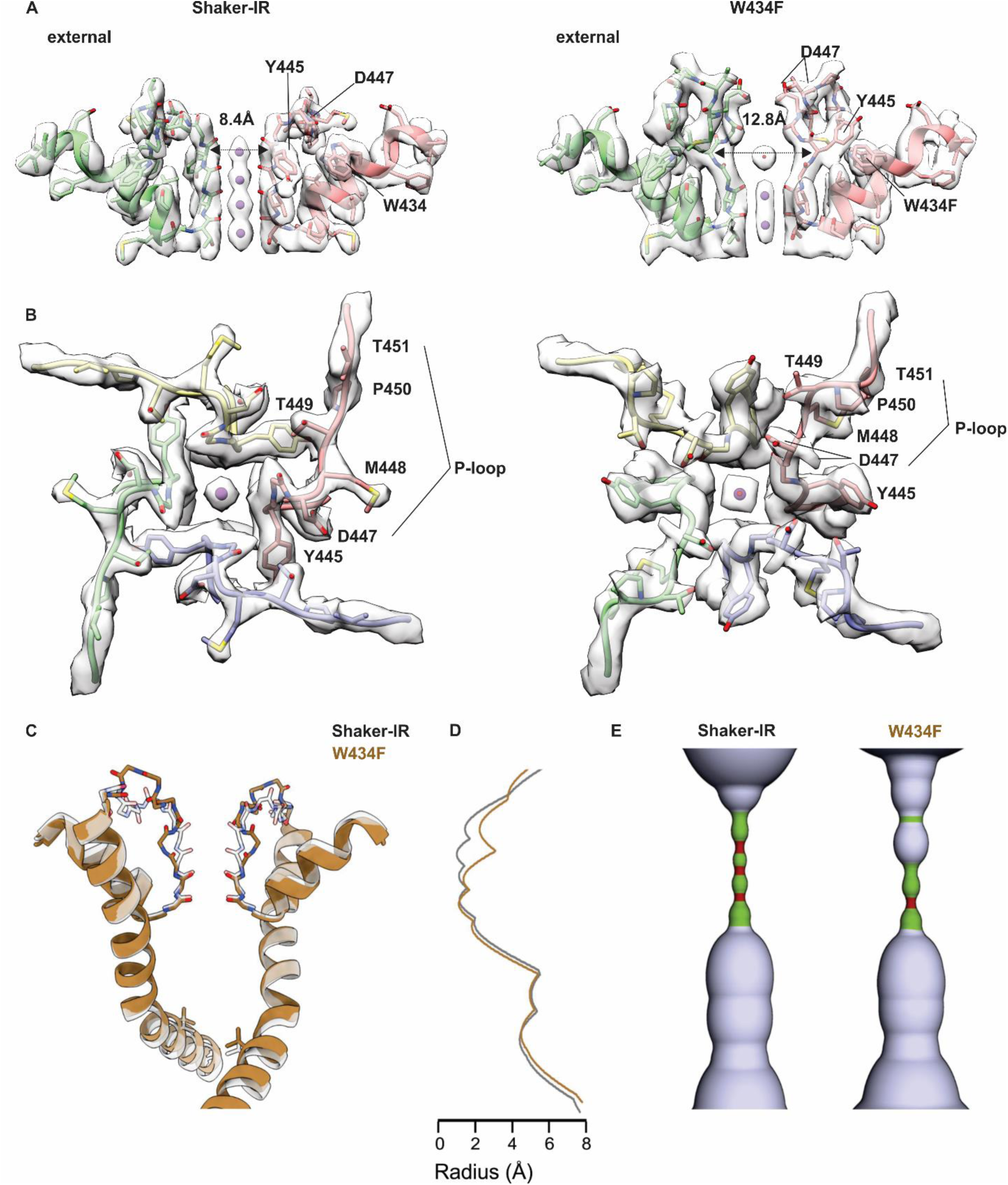
Structure of the external pore of wild-type Shaker-IR and the W434F mutant. **A**, Side-view of the selectivity filter of Shaker-IR (left) and W434F (right). The distance between Y445 CA is shown to highlight the dilation of the selectivity filter in W434F. **B**, View of the pore domain of Shaker-IR (left) and W434F (right) viewed from an extracellular perspective. Residues experiencing a large movement are labeled to highlight structural reorientation of the outer pore domain in W434F. **C**, Superposition of pore-lining regions for Shaker-IR and the W434F mutant. V478 is shown to mark the position of the internal S6 gate that prevents ion permeation across the internal pore when the Shaker Kv channel closes (*51*). **D**, Plot of pore radius for Shaker-IR and the W434F mutant. **E**, Hole diagrams illustrating the ion permeation pathways for Shaker-IR and W434F. Radii ≤ 1 Å are shown in red, radii > 1 Å and ≤ 2 Å are shown in green, and radii larger than 2 Å are shown in light blue. The plot in b and the diagram in c are aligned with the protein model in a.

Within the external pore, in the ion selectivity filter, backbone carbonyl oxygens line the ion permeation pathway and are positioned similarly to other K^+^ channel structures thought to represent a conducting state (*52*) (**Figure 2A**). Strong EM densities can be clearly discerned at four positions within the filter and along its central axis (**Figure 2A**); these densities are consistent with those revealed in previous structural studies (*52*), and appear to reflect a series of ion binding sites within the filter. In summary, there is every indication that the structure of Shaker-IR we have resolved represents an activated, open and conducting conformation.

### Structure of the W434F mutant Shaker Kv channel in lipid nanodiscs

The W434F mutant of Shaker-IR has been extensively characterized and shown to be effectively non-conducting because the mutation accelerates entry of channels into the C-type inactivated state upon membrane depolarization (*24, 30, 41, 42*). We introduced the W434F mutation into our structural construct, expressed and purified the protein and reconstituted it into lipid nanodiscs using the same approach as for Shaker-IR (**Fig. S1**). We solved the structure of the W434F mutant with an overall resolution of 2.9 Å within the TM region; as for Shaker-IR, the cryo-EM map is of excellent quality throughout, enabling us to model the complete sequence of the TM region with the only exception of two external loops in the voltage-sensors (**Fig. S3, S6 and Table S1**). Structural alignment of the S1-S6 regions of the W434F mutant and Shaker-IR reveals their backbone folds are very similar (**Fig. S7**), with low Cα RMSD (1.07 Å) and comparably high TM score (0.9). The structures of the S1-S4 voltage-sensing domains in the two proteins are essentially indistinguishable, and in both instances the internal pores are clearly open **(Figure 2C-E; Fig. S7**).

In contrast, a pronounced conformational change occurs within the external pore (**Figure 2A,B; Video S1**), a region where both cryo-EM maps have the highest resolution (**Extended Figs, 2,6**). A key interaction between D447 in the P-loop and W434 in the pore helix, observed in Shaker-IR and most other K^+^ channels (*7, 41, 52*), is broken in W434F, resulting in a displacement of the P-loop towards the external solution by about 5 Å (**Figure 2A,B; Figure 3A; Fig. S7B**). At the external end of the ion selectivity filter, the sidechain of Y445 undergoes a drastic reorientation in the mutant, breaking off its hydrogen-bond contact with T439 in the adjacent subunit and rotating 90° to reposition behind the filter of the same subunit (**Figure 2A,B; Figure 3A; Fig. S7B**). M448 and T449, two critical positions for C-type inactivation (*25, 28, 32*), also move large distances, with the Cα of M448 moving about 5 Å from an exposed position in Shaker-IR to a buried site near to W435 in the rapidly inactivating mutant (**Figure 3A; Fig. S7B**). In contrast, both T449 and P450 flip from being partially buried in Shaker-IR to completely exposed to solvent in W434F (**Figure 2A,B; Figure 3A; Fig. S7B,C**). Interestingly, the cryo-EM density map for the mutant indicates two distinct rotamers of the D447 side chain, one of which positions the acidic residue near T449 (**Figure 2B; Figure 3A; Fig.S7C**). It is apparent, therefore, that there is an overall reorganization of side chain interactions behind the filter in the inactivated state, with critical residues within the P-loop between Y445 and P450 undergoing substantial displacements. Importantly, this reorganization translates into a marked expansion at the external portion of the ion selectivity filter, by about 4 Å (**Figure 2A-E; Video S1**). Ion densities therein are perturbed, while those at the internal end of the filter are seemingly unaltered (**Figure 2A; Fig. S8**). Inspection of maps and half maps with either C4 or C1 symmetry imposed shows the density within the dilatated portion of the filter is relatively weak, suggestive of occupancy by water molecules or possibly K^+^ ions (**Figure 2A; Fig. S8**).

**Figure 3.**
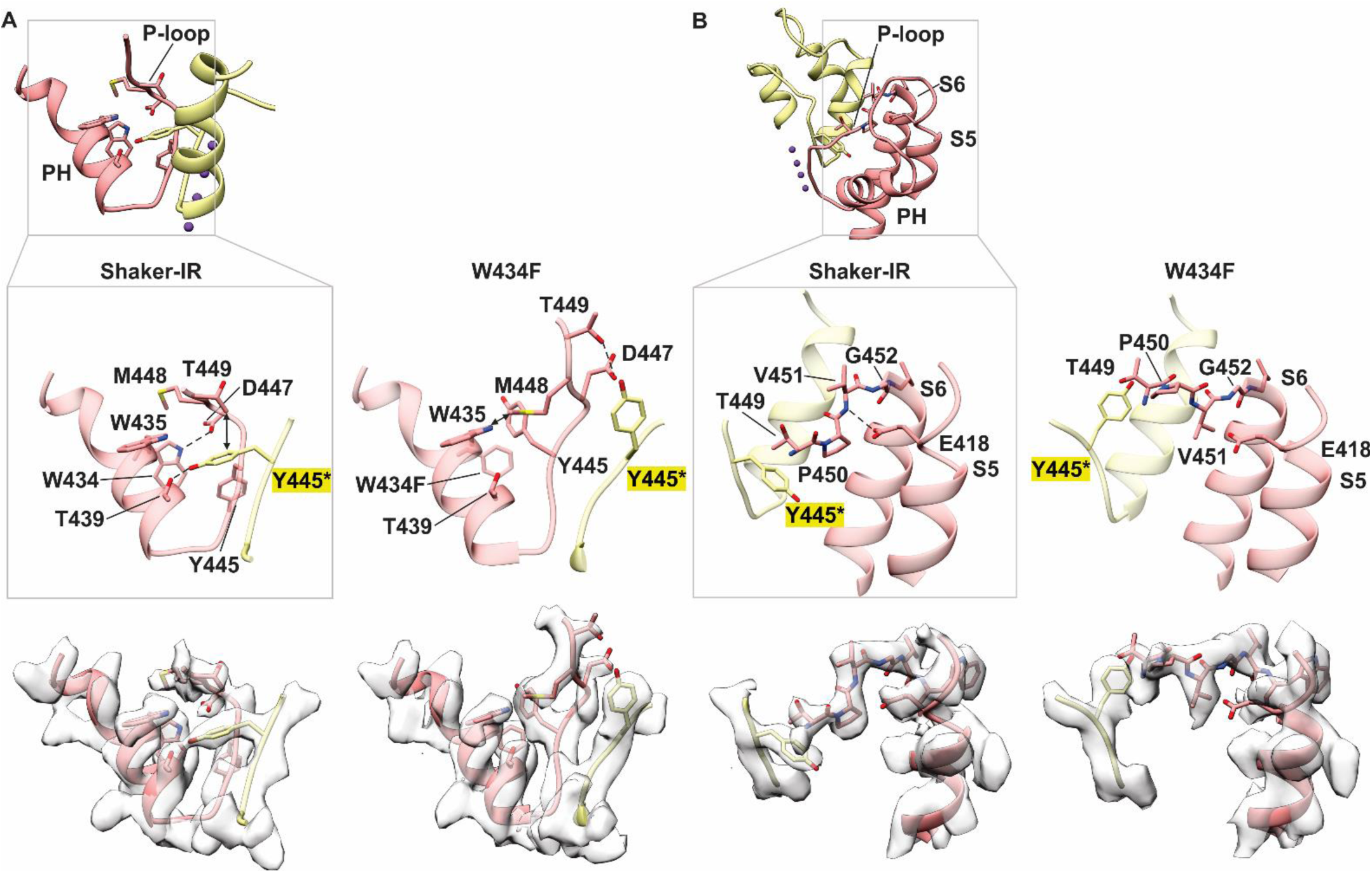
Conformational changes reposition residues that are critical for C-type inactivation in Shaker Kv channels. **A**, Residue interactions within the pore helix (PH) and P-loop for Shaker-IR and the W434F mutant. Inset above provides the orientation of PH and P-loop for two subunits. Enlarged models are shown without or with cryo-EM density (below). D447 forms a hydrogen bond with W434 (OE – NE, 3.3 Å) in Shaker-IR but not in W434F. CG of T449 is 3.6 Å from Y445 in the adjacent subunit in Shaker-IR but 7.5 Å in W434F given movement of both residues. T439 forms a hydrogen bond with Y445 (2.4 Å) in the adjacent subunit in Shaker-IR, which is broken in W434F given the large movement of Y445. In W434F, OG of T449 is 3.2 Å from the OE of D447 and the side chain of M448 is 3.5 Å from W435. **B**, Residue interactions between E418 in S5 with residues in the P-loop for Shaker-IR and the W434F mutant. Inset above provides the orientation of S5, PH, P-loop and S6 for two subunits. Enlarged models are shown without or with cryo-EM density (below). E418 at the top of S5 forms a hydrogen bond with the backbone N of V451 in the P-loop (3.1 Å) in Shaker-IR, but not in W434F (4.3 Å). The pronounced flipping of the side chain of V451 in the W434F mutant directs its side chain towards E418.

It seems apparent that the balance of ion-protein and ion-solvent interactions that controls K^+^ permeation in the activated state is somehow perturbed by the dilation of the external portion of the filter (**Figure 2C-E**) and the reconfiguration of key sidechains such as D447. However, whether conduction is indeed impaired in W434F is not self-evident from the structure alone. To evaluate this question, we calculated a series of molecular dynamics (MD) trajectories specifically designed to evaluate the ion conducting properties of the pore domain in each of the two structures determined experimentally (**Materials and Methods, Fig. S9**). For Shaker-IR, we calculated a trajectory of 3.5 µs that included a transmembrane voltage jump from 0 to 300 mV at t = 0.5 µs. In symmetric 300 mM KCl, we observed a total of 12 complete permeation events after the voltage jump (**Figure 4A,C,D**). In all cases the K^+^ ions traversed the filter in the outward direction and in a stepwise manner, transiently interacting with each of the carbonyl groups along the way (**Movie S2**). One occupancy state however emerges as the most populated at 300 mV (and 300 mM KCl), featuring three K^+^ ions: at the external end of the filter, one K^+^ is in part coordinated by the carbonyls of Y445 and in part hydrated; a second ion concurrently interacts with the carbonyls of G444 and V443; and the third is approximately at same level as the T442 carbonyls (**Figure 4C,D**). As has been noted for other channels, K^+^ permeation proceeds through a knock-on mechanism (*52–54*); interestingly, we observe that this mechanism is at times mediated by water molecules permeating in between adjacent K^+^ ions, while at other times the K^+^ ions interact directly (**Figure 4A,C, Movie S2**). It is also worth noting that the single-channel conductance inferred from this trajectory is approximately 3.5 pS, or 10-fold slower than the value experimentally measured in symmetric 300 mM KCl (*55*). Given the myriad approximations and simplifications adopted in MD simulations, this discrepancy appears reasonable, and consistent with existing literature (*56*). Errors in the order of 1 kcal/mol in the rate-limiting barriers for conduction are most certainly expected and would be sufficient to explain the observed discrepancy. Thus, while additional simulations will be no doubt of interest from both mechanistic and methodological standpoints, we posit that the existing data reaffirms the notion that the structure of Shaker-IR reported here captures the conductive form of the channel.

**Figure 4.**
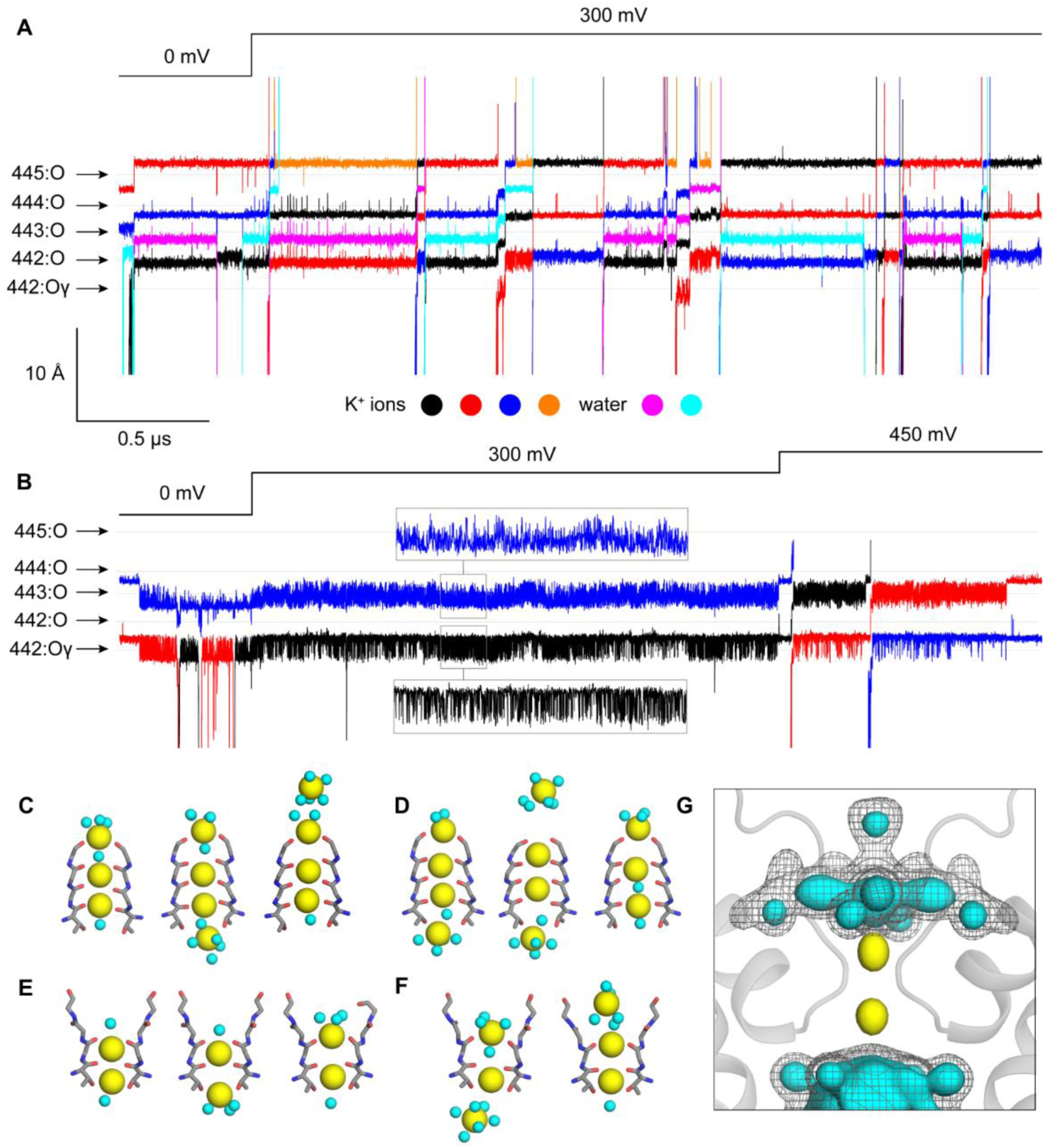
Altered ion interactions with the filter during C-type inactivation in Shaker Kv channels. **A**, Molecular dynamics simulation of K^+^ permeation across the pore domain of Shaker-IR. The figure shows time-series of the position of K^+^ ions as they traverse or transiently enter the selectivity filter; time-series of water molecules traversing the filter are also shown. The time resolution is 120 ps. The relative position of coordinating backbone/sidechain oxygens in residues 442-445 is indicated. Note no voltage was applied during the first 0.5 µs of the trajectory; a jump to 300 mV was introduced at this point and sustained for 3 µs thereafter. **B**, Same as c, for W434F mutant. Note that in this case a second voltage jump from 300 to 450 mV was introduced at 2.5 µs and sustained for 1 µs thereafter. **C,D**, Snapshots of two K^+^ permeation events during the simulation of Shaker-IR. K^+^ ions and water molecules are shown as yellow and cyan spheres, respectively. The elapsed times between snapshots are approximately 3 and 4 ns, respectively. **E**, Representative configurations of the selectivity filter in the simulation of the W434F mutant at 300 mV. Both ions fluctuate but fail to permeate in 2 µs of trajectory under this condition. **F**, Snapshots of a K^+^ permeation event during the simulation of W434F, after the jump to 450 mV. The elapsed time between snapshots is approximately 8 ns. **G**, Calculated iso-density maps for K^+^ (yellow surface) and water (cyan surface, gray mesh) near and within the selectivity filter of W434F, based on analysis of 2 µs of trajectory under 300 mV. For water, two iso-values are used, to illustrate the emergence of well-defined, 4-fold symmetric structures resulting from interactions with ions and protein as well as confinement.

The simulation calculated for the W434F mutant, which was designed identically to that carried out for Shaker-IR and is equivalent in total length, shows radically different results. We did not observe any permeation events before or after the voltage jump from 0 to 300 mV for a total of 2.5 µs of simulation (**Figure 4B, Movie S3**). Instead, two K^+^ ions persistently reside in the filter, coordinated by the carbonyls of V443 and the sidechain and backbone of T442 (**Figure 4E**), while multiple water molecules occupy the pocket made available near G444 and Y445 upon dilation of the filter (**Figure 4G**). The occupancy of water molecules within the external end of the filter in the simulations suggests that the weak density observed within the cryo-EM maps in this region (**Figure 2A;** Fig. S8) is more likely to be water than a K^+^ ion. These observations notwithstanding, we reasoned that because the selectivity filter of W434F is dilated, and not constricted, conduction ought to be feasible without a structural change provided a sufficiently strong driving force. Indeed, a second voltage jump from 300 to 450 mV during the last 1 µs of the trajectory showed 2 full permeation events (**Figure 4B,F**). The permeation mechanism of W434F differs from that observed for Shaker-IR, however, not only in the number of K^+^ ions involved, but also in that K^+^ ions do not dwell at the dilation site near G444 and Y445 before reaching the external solution (**Figure 4B, Movie S3**). Analogous voltage jumps from 300 to 600 mV and from 300 to 900 mV led to a larger number of K^+^ ion permeation events, but nevertheless the preferred species at this distortion was again water (not shown). From this data, we can extract two tentative conclusions: first, the reconfiguration the external pore observed in the W434F mutant severely impairs the conducting properties of the selectivity filter in comparison to Shaker-IR at the same conditions; second, this reconfiguration impairs conduction not by sterically blocking the ion pathway but rather by imposing a mechanism of K^+^ permeation that entails a significantly larger energetic barrier, as a result of the filter dilation. Whether K^+^ ions can traverse this barrier at all in experimentally accessible conditions remains however an open question. It will be of great interest to further dissect and quantify these phenomena in future studies.

## Discussion

Our objective was to solve structures of the Shaker Kv channel to understand its mechanism of C-type inactivation, which is likely shared by many other eukaryotic channels. The structure of Shaker-IR provides an important framework for future studies aimed at exploring voltage-dependent gating mechanisms, and for reexamining a large body of functional studies to gain new mechanistic insights (*1, 4, 5*). The value of structures of the same protein used in extensive functional studies is exemplified here with the W434F mutation, which renders the channel effectively non-conducting by greatly promoting C-type inactivation (*24, 30, 42*).

Is the conformational rearrangement and dilation of the selectivity filter that we observe in W434F representative of the structural change occurring during C-type inactivation in wild-type Shaker Kv channels? Although additional studies are warranted explore this question further, and C-type inactivation might involve multiple distinct conformational changes in the outer pore, the structural changes we observe in W434F are remarkably consistent with many functional studies exploring the mechanism of C-type inactivation in the Shaker Kv channel. The hydrogen bond between D447 and W434 that breaks off in the structure of W434F (**Figure 3A**) has been shown to stabilize the open state of the Shaker Kv channel (*41*), with mutation of either residue dramatically promoting C-type inactivation (*24, 30, 32, 41, 57*). T449 is a particularly critical residue for C-type inactivation in the Shaker Kv channel because polar substitutions speed C-type inactivation and hydrophobic substitutions slow inactivation (*25*). Indeed, hydrophobic substitutions at T449 so dramatically slow C-type inactivation that they can rescue normal ion conduction in W434F (*29*). The structure of Shaker-IR reveals that T449 is partially buried as it is positioned within 4 Å of the aromatic ring of Y445 in the adjacent subunit, whereas it becomes completely solvent exposed in the C-type inactive state (**Figure 3A**). Hydrophobic substitution at 449 would thus be expected to stabilize the conducting conformation by strengthening hydrophobic interactions between 449 and Y445. Remarkably, the large displacement of T449 in W434F places this sidechain within 3.2 Å of D447 for one of the resolved rotamers for the D447 sidechain (**Figure 3A**), which explains why Cys substitutions at position 449 generate a strong metal binding site that forms at much faster rates when Shaker Kv channels are inactivated (*26*). Although these metal bridges had been envisioned to form between T449C residues in opposing subunits across the pore axis, thereby supporting the theory that C-type inactivation entails the collapse of the filter, our structures reveal an alternate interpretation, namely that the metal bridges form between T449C and D447 within the same subunit in the C-type inactivated state. Indeed, external Ba^2+^ ions have been shown to inhibit the Shaker Kv channel at a site that is sensitive to substitutions at both T449 and D447 (*57*), which can be readily understood given the intimate relationship between these residues seen in the W434F structure. The increased solvent exposure of T449 and P450 observed in the W434F structure is also consistent with the enhanced rate of modification of introduced Cys residues at these positions by thiol reactive compounds when Shaker C-type inactivates (*28*). The intersubunit hydrogen bond between T439 and Y445 observed in Shaker-IR and that is necessarily broken with the large movement of Y445 that occurs in the structure of W434F (**Figure 3A**), has also been shown to stabilize the open state of Shaker Kv channels, with mutation of either residue dramatically promoting C-type inactivation (*41*). Finally, the structural changes we observe can explain a series of important observations around E418 within the external end of the S5 helix near the perimeter of the pore domain (**Figure 3B**). Introduction of fluorophores at position S424 in the Shaker Kv channel, which in the structure of Shaker-IR is nearby to E418, give rise to fluorescence changes that closely track C-type inactivation (*58*). Mutations of E418 dramatically speed C-type inactivation in Shaker Kv channels (*59, 60*), and substitutions with Cys at E418 and V451 within the P-loop lead to disulfide bond formation that stabilizes the C-type inactivated state (*60*). In the structure of Shaker-IR, E418 hydrogen bonds with the backbone amide of V451 in the P-loop and this interaction is lost as the P-loop is displaced in W434F (**Figure 3B**). Notably, displacement of the P-loop in the W434F mutant is accompanied by flipping of the side chain of V451 towards E418 (**Figure 3B**). Although side chain density is missing for E418 in the W434F mutant, the EM density for the external end of the S5 helix is similar in Shaker-IR and W434F (**Figure 3B**), consistent with E418C and V451C being optimally positioned to form a disulfide bond that stabilizes the C-type inactivated state.

It is important to clarify that our structures are not at all inconsistent with the observation that externally-applied tetraethyl ammonium or K^+^ ions can interfere with C-type inactivation in Shaker Kv channels. While this observation had been rationalized using a foot-in-the-door analogy, in support of the notion the filter collapses upon inactivation (*26, 28*), a mechanism whereby binding of these ions within the selectivity filter hinders inactivation by precluding its dilation is equally plausible (*15*). All that is required to explain this particular experimental observation is for the conducting filter to interact more favorably with external ions than the inactive state; the structure of the W434F mutant of the Shaker Kv channel is clearly consistent with this premise, as it lacks the external ion-binding sites seen in the activated state.

The structural changes we observe in the ion selectivity filter also provide important insight into how inactivation of the selectivity filter might be coupled with other conformational mechanisms elsewhere in the channel. For example, in KcsA opening of the internal pore is thought to trigger collapse of the selectivity filter because a bulky Phe in the pore-lining M2 helix interacts with a conserved Thr at the base of the ion selectivity filter (**Fig. S10D**) (*38, 61*). Our structures suggest that this coupling mechanism is unlikely to apply for the Shaker Kv channel, as the base of the ion selectivity filter is unchanged when comparing Shaker-IR and the W434F mutant (**Figure 3; Fig. S10C**). The reason for this difference might be that the residue equivalent to the bulky Phe in the S6 helix of Shaker is I470 (**Fig. S10C**), or it might simply be that the conformational changes during inactivation in KcsA are unrelated to those occurring in Shaker (**Fig. S10**). Indeed, C-type inactivation in Shaker is thought to be coupled to the conformation of the voltage-sensing domains (*58, 60, 62*). In the future it will be interesting to further explore coupling between voltage-sensor activation, internal pore opening and C-type inactivation by solving structures of the Shaker Kv channel with voltage sensors in resting states and the pore in a closed state.

A structural mechanism for C-type inactivation through filter dilation raises the interesting question about what conditions might promote a more constricted or collapsed conformation, like that observed in the KcsA channel in the presence of low K^+^ concentrations (*52*). If a K^+^-dependent collapse of the filter also occurs in Kv channels, it would likely represent a conformation adopted each time the internal gate of a K^+^ channel closes under physiological ionic conditions because in that state the filter will ultimately equilibrate with the external solution containing low [K^+^] (**Figure 5**). Collapse of the filter would assure that ion permeation is minimized when the internal gate closes in the absence of an activating stimulus (*46*). It will be interesting to determine whether the ion selectivity filters in Kv channels collapse in low K^+^ because it would indicate that the filter adopts at least three distinct conformations during the gating cycle of a Kv channel. It will also be exciting to determine whether other K^+^ channels use filter dilation to regulate ion permeation. Although structural changes related to those we see here have thus far not been resolved within the filter of K2P channels, for example, closure of those channels has been proposed to involve rearrangement of residues in the P-loop and dilation of the filter and has been shown to deplete ion occupancy in the external half of the filter while maintaining those within the inner half (*63*), suggesting that the mechanisms of C-type inactivation in Shaker and filter gating in K2P channels may be related.

**Figure 5.**
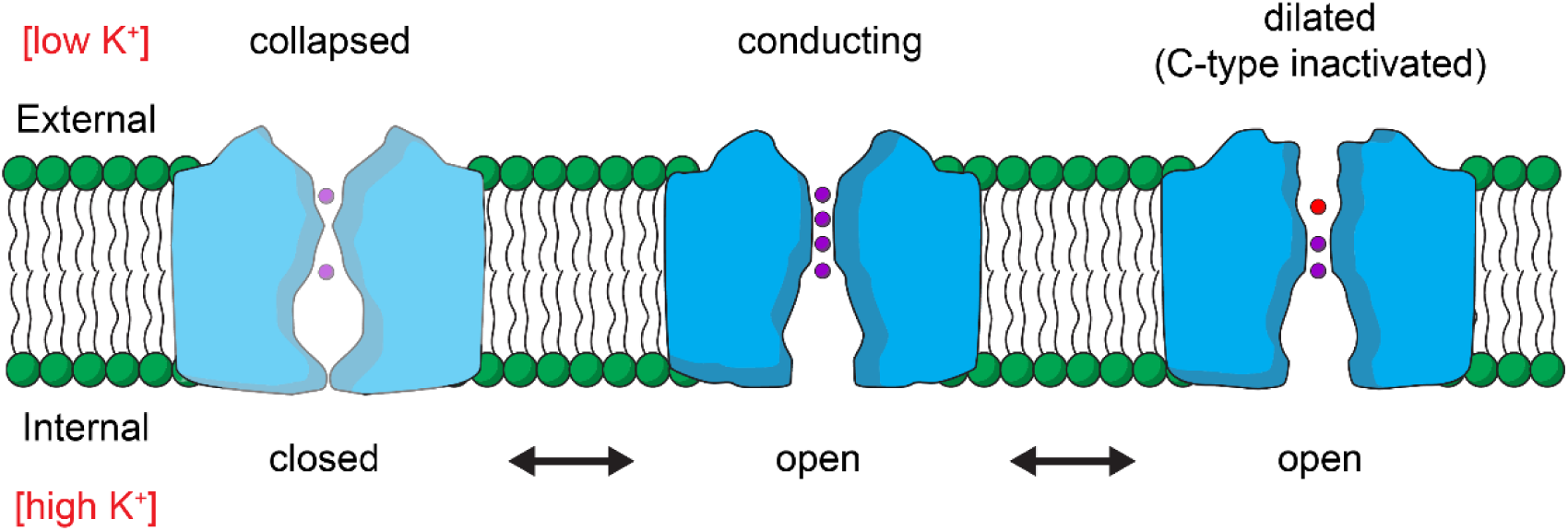
Gating cycle of the pore domain of a Kv channel. Cartoons illustrating dynamic changes in structure of the ion conduction pore during the gating cycle of a Kv channel. With voltage-sensor activation and opening of the internal S6 gate, the filter will equilibrate with high potassium inside the cell and adopt a conducting conformation. With prolonged activation, the filter C-type inactivates and dilates to diminish ion permeation. Four K^+^ ion binding sites (purple) within the filter can be occupied in the conducting state, whereas in the C-type inactivated state ions primarily occupy only the internal half of the filter and water molecules (red) likely occupies the dilated external part of the filter. At negative membrane voltages where the voltage-sensing domains are resting and the internal S6 gate is closed, the filter would equilibrate with the external solution containing low potassium and potentially collapses, as proposed based on studies of the KcsA potassium channel (*52*).

## Materials and Methods

### Shaker Kv channel expression using Baculovirus and mammalian expression system

To produce the Shaker Kv channel for cryoEM, WT and W434F channels were cloned into the pEG vector in which EGFP was substituted with mVenus (*64*) and expressed in tsA201 cells using the previously published Baculovirus-mammalian expression system with a few minor modifications (*65*) . Briefly, P1 virus was generated by transfecting Sf9 cells (∼2.5 million cells on a T25 flask with a vent cap) with 50 – 100 ng of fresh Bacmid using Cellfectin. After 4 – 5 days incubation in a humidified 28 °C incubator, the cell culture media was collected by centrifugation (3,000g x 10 min), supplemented with 2% FBS, and filtered through a 0.45 μm filter to harvest the P1 virus. To amplify the P1 virus, ∼500 ml Sf9 cell cultures at a ∼1.5 million cells/ml density were infected with 1 – 200 μl of the virus and incubated in a 28 °C shaking incubator for 3 days. The cell culture media was then collected by centrifugation (5,000g x 20 min), supplemented with 2% FBS, and filter through 0.45 μm filter to harvest P2 virus. The volume of P1 virus used for the amplification was determined by carrying out a small-scale amplification screening in which ∼10 ml Sf9 cell cultures at the same density were infected with different volume of P1 virus and harvested after 3 days to transduce tsA201 cells and compare the expression level of Shaker Kv channels using mVenus fluorescence intensity. The P2 virus was protected from light using aluminum foil and stored at 4 °C until use. To express the Shaker channels, tsA201 cells at ∼1.5 million cells/ml in Freestyle medium with 2 % FBS were transduced with 10 % (v/v) P2 virus and incubated at a 37 °C CO_2_ incubator. To boost the protein expression, sodium butyrate (2 M stock in H_2_O) was added to 10 mM at ∼16 hours of post-transduction. The culture was continued at the 37 °C CO_2_ incubator for another 24 hours, and the cells were harvested by centrifugation (5,000g x 20 min) and frozen at -80 °C until use.

### Shaker Kv channel purification

Prior to extraction of the Shaker channels from tsA201 cells, membrane fractionation was carried out using a hypotonic solution and ultracentrifugation. The cells were first resuspended in a hypotonic solution (20 mM Tris pH 7.5 and 10 mM KCl) with protease inhibitors (pepstatin, aprotinin, leupeptin, benzamidine, trypsin inhibitor, PMFS) using a Dounce homogenizer, incubated at 4 °C for ∼30 minutes, and centrifuged at 1,000 g for 10 minutes to remove cell debris.

The supernatant was ultracentrifuged for 1 hour (45,000 rpm, Beckman Ti45 rotor) and collected membranes were stored at -80 °C until use. To purify Shaker Kv channels, the fractionated membranes were resuspended in an extraction buffer (50 mM Tris pH 7.5, 150 mM KCl, 2 mM TCEP, 50 mM DDM, 5 mM CHS with the protease inhibitor mixture used above) and extracted for 1 hour at 4 °C. The solution was clarified by centrifugation (12,000g x 10 min) and incubated with CoTALON resins at 4 °C for 1 hour, at which point the mixture was transferred to an empty disposable column (Econo-Pac® Biorad). The resin was washed with 10 column volume of Buffer A (50 mM Tris pH 7.5, 150 mM KCl, 1 mM DDM, 0.1 mM CHS, and 0.1 mg/ml porcine brain total lipid extract) with 10 mM imidazole, and bound proteins were eluted with Buffer A with 250 mM imidazole. The eluate was concentrated using Amicon Ultra (100kDa) to ∼350 – 450 μl and loaded onto a Superose6 (10x300mm) gel filtration column and separated with Buffer A. All purification steps described above was carried out at 4 °C or on ice.

### Lipid nanodisc reconstitution of the Shaker Kv channel

Lipid nanodisc reconstitution was performed following the previously published methods with minor modifications (*8*) . On the day of nanodisc reconstitution, the Shaker Kv channel purified by Superose6 in detergent was concentrated to ∼1–3 mg/ml and incubated with histidine tagged MSP1E3D1 and 3:1:1 mixture of 1-palmitoyl-2-oleoyl-sn-glycero-3-phosphocholine (POPC), 1-palmitoyl-2-oleoyl-sn-glycero-3-phospho-(1’-rac-glycerol) (POPG) and 1-palmitoyl-2-oleoyl-sn-glycero-3-phosphoethanolamine (POPE) for 30 minutes at room temperature. The mixture was transferred to a tube with Biobeads (∼30–50 fold of detergent; w/w) and incubated at room temperature for ∼3 hours in the presence of TEV protease (prepared in-house) and 2 mM TCEP to remove N-terminal fusion protein including poly-histidine and mVenus tag. The reconstituted protein was loaded onto Superose6 column (10x300mm) and separated using 20 mM Tris and 150 mM KCl buffer at 4 °C. The success of nanodisc reconstitution was confirmed by collecting separated fractions and running SDS-PAGE to verify the presence of Shaker Kv and MSP1E3D1 bands at a similar ratio. Typically, optimal reconstitution required the incubation of 1:10:200 or 1:10:400 molar ratio of tetrameric Shaker Kv, MSP1E3D1, and the lipid mixture.

### Cryo-EM sample preparation and data acquisition

Concentrated samples of Shaker-IR or the W434F mutant in nanodiscs (3 µL) were applied to glow-discharged Quantifoil grids (R 1.2/1.3 Cu 300 mesh). The grids were blotted for 2.5 s, blot-force 4, 100% humidity, at 16 °C using a FEI Vitrobot Mark IV (Thermo Fisher), followed by plunging into liquid ethane cooled by liquid nitrogen. Images were acquired using an FEI Titan Krios equipped with a Gatan LS image energy Filter (slit width 20eV) operating at 300 kV. A Gatan K3 Summit direct electron detector was used to record movies in super-resolution mode with a nominal magnification of 105,000x, resulting in a calibrated pixel size of 0.43 Å per pixel. The typical defocus values ranged from – 0.5 to -1.5 um. Exposures of 1.6 s were dose-fractionated into 32 frames, resulting in a total dose of 52 e- Å-2. Images were recorded using the automated acquisition program SerialEM (*66*).

### Image processing

All processing was completed in RELION (*67*) and cryoSPARC (*68*). The beam-induced image motion between frames of each dose-fractionated micrograph was corrected and binned by 2 using MotionCor2 (*69*) and contrast transfer function (CTF) estimation was performed using CTFFIND4 (*70*). Micrographs were selected and those with outliers in defocus value and astigmatism, as well as low resolution (>5 Å) reported by CTFFIND4 were removed. The initial set of particles from 300 micrographs were picked using Gautomatch (https://www2.mrc-lmb.cam.ac.uk/research/locally-developed-software/zhang-software/#gauto) and followed by reference-free 2D Classification in RELION. The good classes were then used as template to pick particles from all selected micrographs using Gautomatch. 4,934,881 (Shaker-IR) and 7,446,104 (W434F) particles were picked and extracted with 2x downscaling (pixel size 1.72 Å). Several rounds of reference-free 2D Classification were performed to remove ice spot, contaminants and bad particles, yielding 2,246,076 (WT) and 2,347,164 (W434F) particles, respectively. The good particles were 3D classified using reference generated by 3D initial model in C1 symmetry. Good classes were then selected and followed by further 2D Classification and 3D Classification. The classes which show good features within the transmembrane domains were merged. After removing duplicate particles, the selected particles were re-extracted using box size 296 pixels without binning (pixel size 0.86 Å). To get good reconstruction within the transmembrane domain, the re-extracted particles were further 3D classified with a transmembrane domain mask in C4 symmetry, followed by 3D auto-refine and CTF refinement. After that, Bayesian polishing was performed, and bad particles were removed from polishing particles using 2D Classification. The selected polishing particles were subjected to 3D auto-refine in RELION and then Non-uniform refinement in cyroSPARC. The final reconstruction was reported at 3.0 Å for Shaker-IR and 2.9 Å for W434F. To confirm whether ion density in the selectivity filter is real, the reference was shifted by 10 pixels in X-direction using relion_image_handler so that the selectivity filter is not in the center axis. C1 symmetry was applied for 3D auto-refine. The ion density without postprocessing is shown in **Fig. S8**.

### Model building and structure refinement

Model building was first carried out by manually fitting the transmembrane domain of Kv1.2-2.1 paddle chimera channel (PDB 6EBM) into the electron microscopy density map using UCSF Chimera (*71*). The model was then manually built in Coot (*72*) and refined using real_space refinement in PHENIX (*73*) with secondary structure and geometry restraints. The final model was evaluated by comprehensive validation in PHENIX. Structural figures were generated using PyMOL (https://pymol.org/2/support.html) and UCSF Chimera.

### Structural Alignments

Alignments of single subunits (**Fig. S7**) were performed using align in PyMOL. Alignments of tetramers (**Fig. S4, S10**) were performed using Fr-TM-Align (*74*). Alignments between KcsA and Shaker (**Fig. S10**) were performed using the pore domain (S5-S6 helices) of all four subunits. Alignments between the Kv 1.2/2.1 paddle chimera and Shaker (**Fig. S4**) were performed using the transmembrane domain (S1-S6 helices) of all four subunits. Pore radii were calculated using the HOLE program (*75*).

### Molecular dynamics simulations of Shaker-IR and the W434F mutant

The simulations of Shaker-IR and the W434F mutant used the cryo-EM structures reported in this study as the starting configuration; in both cases, the simulations examined a construct encompassing residues 382 to 485 (from S4-S5 helix to S6 helix), with neutral N- and C-termini and all ionizable sidechains in their default protonation state at pH 7 (the net charge of these constructs is thus zero). For Shaker-IR, two K^+^ ions were initially positioned in the selectivity filter, one coordinated by residues 442 and 443 and another by residues 444 and 445; a third ion was positioned below the sidechain of 442, and two water molecules were modeled between the three ions. For the W434F mutant, K^+^ ions were initially positioned between residues 443 and 444 and by residue 442, with a water molecule in between. (Note neither configuration was ultimately the most populated in the resulting trajectories.) Dowser (*76*) was used to model water molecules buried within the protein structures. To complete the initial set-up of the experimental cryo-EM structures, both constructs (including ions and water molecules) were briefly energy-minimized using CHARMM(*77*) and the CHARMM36m forcefield (*78–80*); specifically, the minimization consisted of 250 steps using the steepest-descent algorithm, followed by 250 steps using the conjugate-gradient algorithm. Both Shaker-IR and W434F were simulated in a POPC lipid bilayer flanked by a 300 mM KCl solution. To generate a molecular model of this membrane/solvent environment, we created a coarse-grained (CG) POPC-lipid bilayer in 300 mM KCl in an orthorhombic box of ∼90 × 90 × 100 Å, using insane.py (*81*). To equilibrate this CG system, we carried out a 20-µs MD simulation using GROMACS 2018.8 (*82*) and the Martini2.2 forcefield (*83–86*), at constant temperature (303 K) and constant semi-isotropic pressure (1 atm) and with periodic boundary conditions. The integration time-step was 20 fs. To embed Shaker-IR in this environment, its atomic structure was first coarse-grained using martinize.py (*83, 85*), and overlaid onto the membrane/solvent system in a configuration that seemed plausible, removing all overlapping lipid/water molecules. To optimize the protein/lipid/solvent interfaces in the resulting model, we carried out a 10-µs MD simulation of the complete system using GROMACS 2018.8 (*82*) and Martini2.2 (*83–86*) at constant temperature (303 K) and constant semi-isotropic pressure (1 atm) and with periodic boundary conditions and an integration time-step of 20 fs. Having ascertained the equilibration of the membrane structure and of the position of the protein in the lipid bilayer, the final snapshot of the CG trajectory was transformed into an all-atom representation based on the CHARMM36m forcefield (*78–80*). To do so, lipid and solvent molecules were back-mapped (*87*) onto all-atom models; and the CG version of Shaker-IR was replaced with the energy-minimized all-atom construct described above, after optimally superposing the Cα trace of the latter onto that of the former. The resulting all-atom molecular system includes 201 POPC lipids, 14,376 water molecules, 79 K^+^ and 79 Cl^-^ (300 mM KCl), in an orthorhombic box of ca. 90 × 90 × 100 Å.

To further optimize this all-atom model, the simulation system was first energy-minimized for 5,000 steps using with NAMD 2.13 (*88, 89*) and the conjugate-gradient algorithm. We then carried out a series of MD simulations wherein structural restraints are applied to the protein and ions/water in the selectivity filter progressively weakened over the course of ∼150 ns. The scheme followed in this multi-step equilibration is summarized in **Table S2**. These simulations were carried out using NAMD 2.13 (*88, 89*) and CHARMM36m (*78–80*) at constant temperature (298 K) and constant semi-isotropic pressure (1 atm) with periodic boundary conditions and an integration time-step of 2 fs. Electrostatic interactions were calculated using PME, with a real-space cut-off value of 12 Å; van der Waals interactions were also cut-off at 12 Å, with a smooth switching function taking effect at 10 Å. The all-atom simulation system for the W434F mutant was constructed by replacing the structure of Shaker-IR with the structure of the W434F mutant at the end of Step 6 in the equilibration protocol summarized in Table 2. The resulting all-atom molecular system includes 201 POPC lipids, 14,375 water molecules, 79 K^+^ and 79 Cl^-^ (300 mM KCl), in an orthorhombic box of ca. 90 × 90 × 100 Å. The complete minimization/equilibration protocol (Steps 1 through 9) was then repeated for the W434F mutant. No external electric field was applied during either the equilibration of Shaker-IR or the W434F mutant, and so the transmembrane voltage in both cases was zero. The dimensions of both simulation systems at the end of the equilibration protocol are ca. 89 × 89 × 92 Å. Finally, to quantify the ion-conducting properties of Shaker-IR and the W434F mutant, a 3.5-µs MD trajectory was calculated for each construct using an Anton 2 supercomputer (*90*) and the CHARMM36m forcefield (*78–80*). The starting configuration for each of these simulations was the final configuration after Step 9 in the corresponding equilibration. These simulations were also carried out at constant temperature (298K) and semi-isotropic pressure (1 atm), respectively set with the Nose-Hoover thermostat (*91, 92*) and the Martyna-Tobias-Klein barostat (*93*), and with periodic boundary conditions and an integration time-step of 2.5 fs. Electrostatic interactions were calculated using the Gaussian split Ewald method (*94*); van der Waals interactions were cut-off at 10 Å. To ensure that the simulations evaluate the properties of the functional states captured by the cryo-EM data, we precluded structural distortions that might occur in the microsecond time-scale by supplementing the forcefield with a series of potential-energy terms that bias but do not confine the dynamics of all Φ, Ψ and χ_1_ dihedral angles towards the values observed in the experimental structures. The functional form of each of these terms is 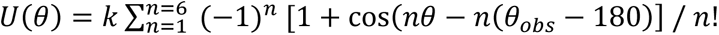, which is identical to that used in previous studies based on microsecond simulations calculated on Anton (*95, 96*), but we used a weaker bias *k* equal to 0.6 kcal/mol. Accordingly, the conformational ensembles in our simulations of Shaker-IR and W434F are quite diverse; for example, typical RMS deviations of the protein backbone from the cryo-EM structures range from 1 to 1.4 Å (**Fig. S9**). No external electric field was applied during the first 0.5 µs of either of MD trajectories. At 0.5 µs, a voltage jump to 300 mV was introduced in the system, for both Shaker-IR and the W434F, through the application of an outwardly directed electric field of 0.075 kcal/mol Å^-1^ e^-1^ perpendicular to the membrane plane (1 kcal/mol Å^-1^ e^-1^ = 43.4 mV/Å). For a W434F, a second voltage jump to 450 mV was introduced at 2.5 µs through the application of an outwardly directed electric field of 0.1125 kcal/mol Å^-1^ e^-1^.

## Acknowledgments

We thank Andres Jara-Oseguera, Miguel Holmgren, Mark Mayer and members of the Swartz laboratory for helpful discussion, and Huaibin Wang in the NIH Multi-Institute Cryo-EM Facility (MICEF) for assistance in acquiring cryo-EM data. This work utilized the NIH MICEF and computational resources of the NIH HPC Biowulf cluster (http://hpc.nih.gov)

## Funding

This research was supported by the Intramural Research Programs of the National Institute of Neurological Disorders and Stroke, NIH, Bethesda, MD to KJS, and the National Heart Blood and Lung Institute, NIH, Bethesda, MD to J.J. and J.F.G.

## Author contributions

Conceptualization: CB, XT, KJS

Methodology: XT, CB, RS, AIFM, KH, TC, JJ, JDF, KJS

Investigation: XT, CB, RS, AIFM, TC, JDF

Visualization: XT, CB, RS, AIFM, KH, JDF, KJS

Funding acquisition: JJ, JDF, KJS

Project administration: CB, JDF, KJS

Supervision: JDF, KJS

Writing – original draft: XT, CB, RS, AIFM, KH, TC, JJ, JDF, KJS

Writing – review & editing: XT, CB, RS, AIFM, KH, TC, JJ, JDF, KJS

## Competing interests

Authors declare that they have no competing interests.

## Data and materials availability

The map of wild-type Shaker-IR and the W434F mutant have been deposited in the Electron Microscopy Data Bank (EMDB) under accession codes EMD-25147 and EMD-25152, respectively. Models of wild-type Shaker-IR and the W434F mutant have been deposited in the Protein Data Bank with accession codes 7SIP and 7SJ1, respectively.

## Supplementary Materials

Figs. S1 to S10

Tables S1 to S2

Captions for Movies S1 to S3

Movies S1 to S3

## Supplementary Materials

**Fig. S1.**
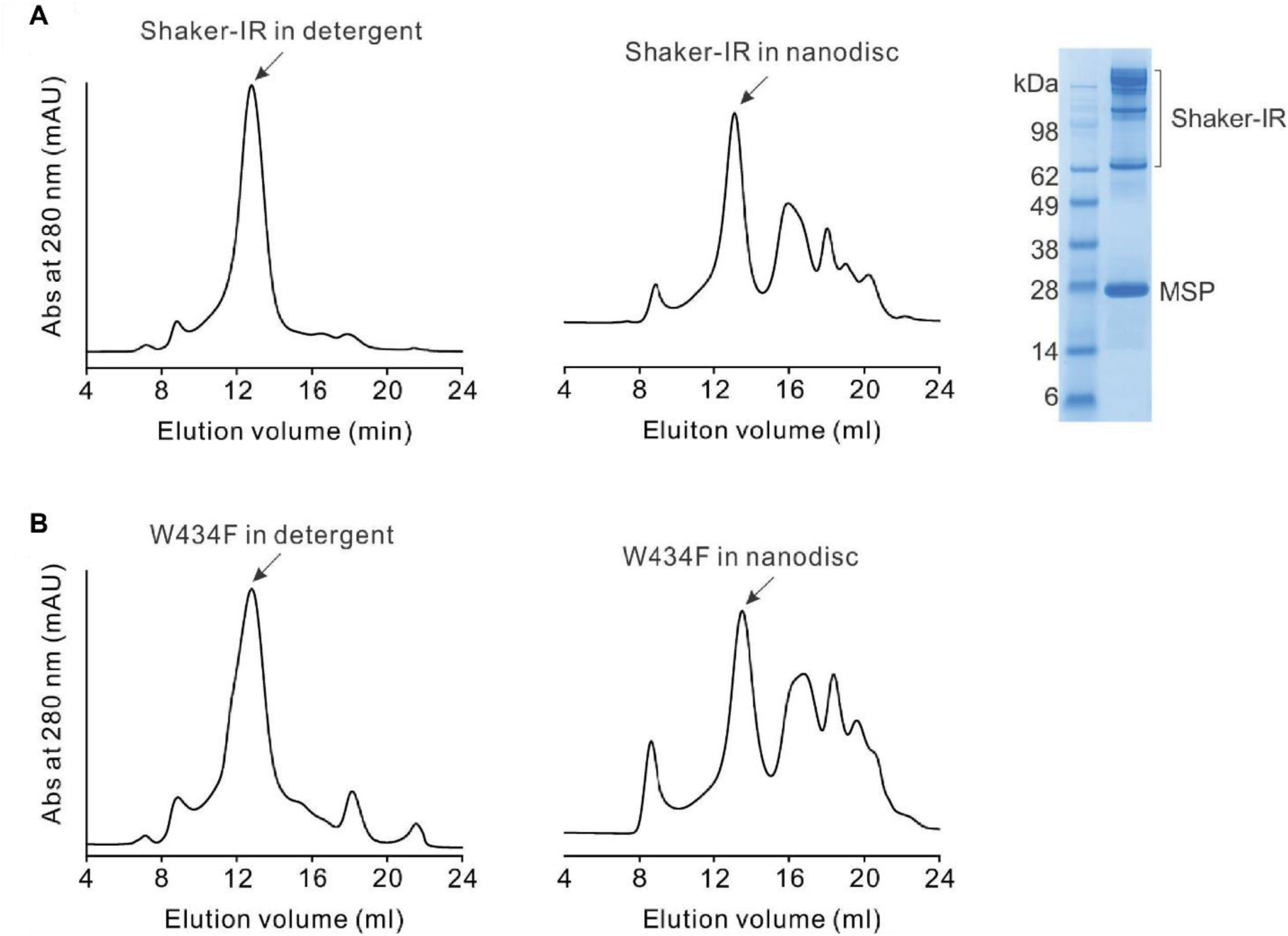
Biochemistry for wild-type Shaker-IR and W434F mutant. **(A)** Gel filtration chromatograms of the WT Shaker in detergent (left) and in nanodisc (middle), and SDS-PAGE of the WT Shaker in nanodisc. The multiple bands running at a higher molecular weight in the SDS-PAGE appear to reflect the glycosylated species of the channel. **(B)** Gel filtration chromatograms of W434F mutant in detergent (left) and in nanodisc (right). The peak fractions marked with the arrow were collected and used either for nanodisc reconstitution or for cryo-EM imaging.

**Fig. S2.**
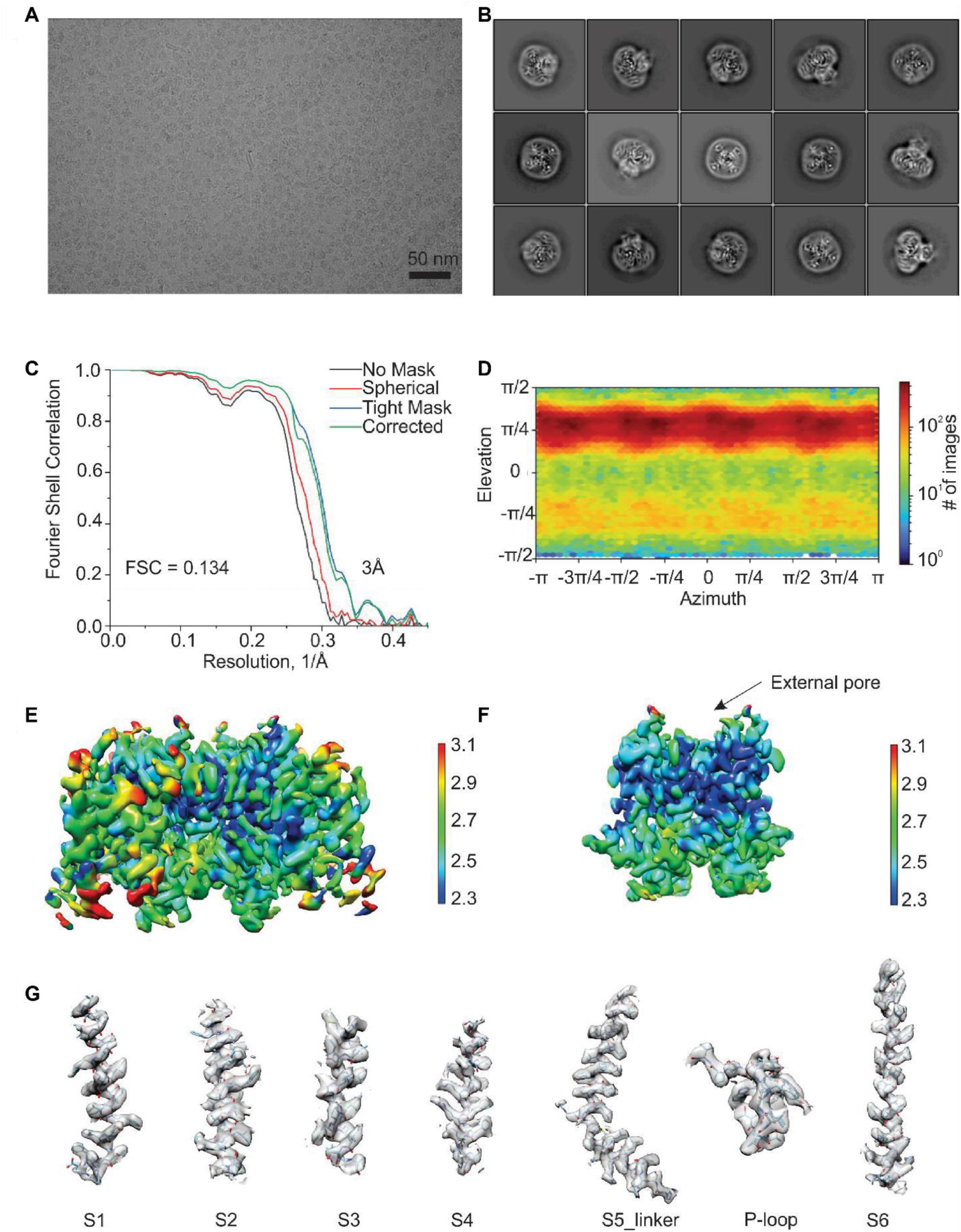
Cryo-EM imaging of Shaker-IR. **(A)** Cryo-EM micrograph of Shaker-IR. **(B)** 2D class averages of the particles in different orientations.**(C)** Fourier Shell Correlation (FSC) curves. **(D)** Direction distribution plots of the 3D reconstruction. **(E)** Local resolution map for the entire TM region. **(F)** Local resolution map for the S5-S6 pore domain, highlighting the dark blue region within the outer pore that shows the best resolution inthe overall structure. **(G)** Regional cryo-EM density for Shaker-IR.

**Fig. S3.**
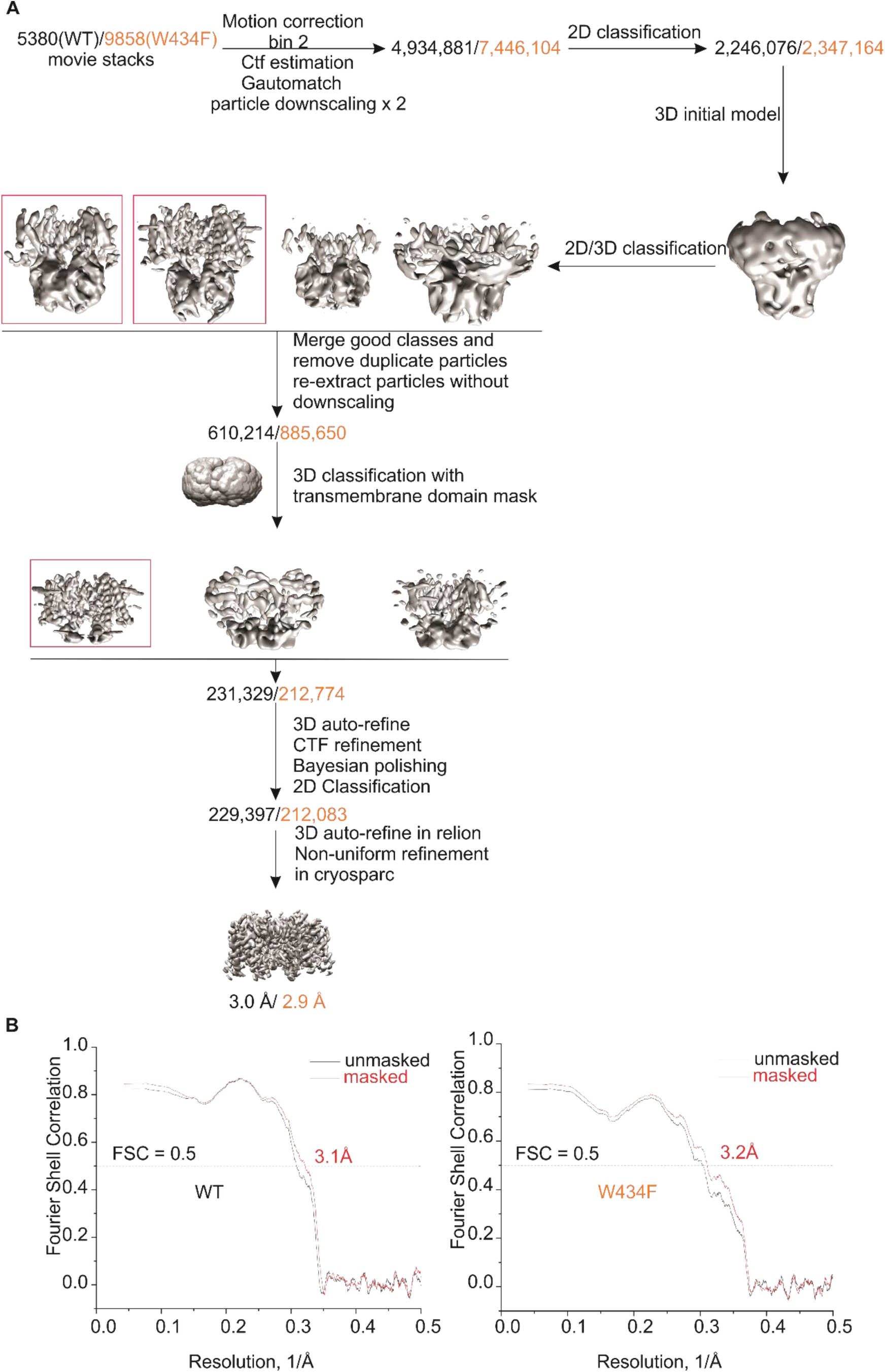
Data processing workflow for the cryo-EM structures of Shaker-IRand the W434 mutant. (A) Workflow for the cryo-EM data processing. (B) FSC curves of the model versus map.

**Fig. S4.**
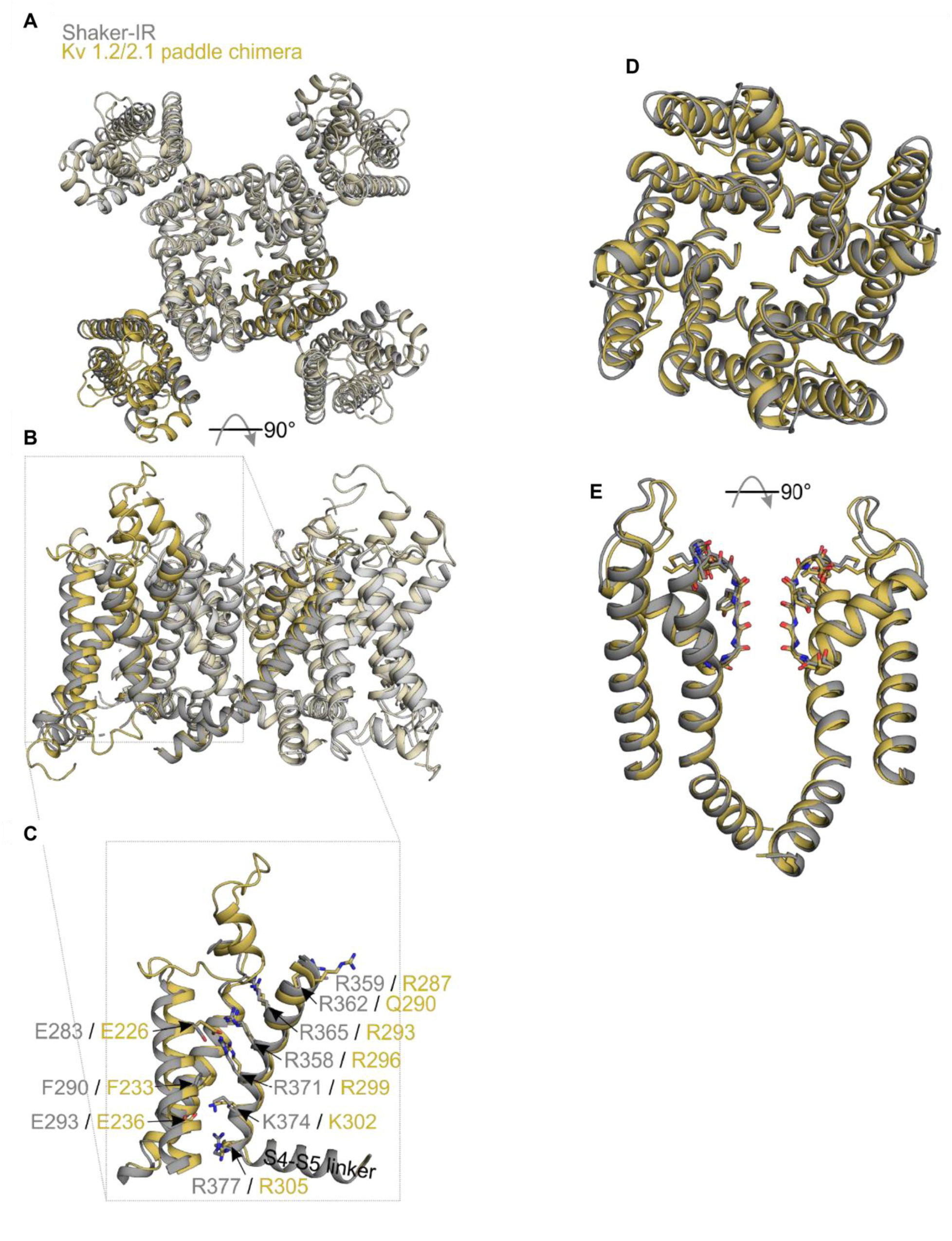
Structural alignment of Shaker Kv channels with the Kv2.1/1.2 paddle chimera. Superimposition of transmembrane domains (S1-S6) of Shaker-IR (7SIP) and the Kv 1.2/2.1 paddle chimera (2R9R) **(A)** top and **(B)** side views, along with **(C)** focused view of one S1-S4 voltage-sensing domain with S4 charges and charge transfer center residue side chains shown as sticks. Superimposition of pore domains (S5-S6) **(D)** top and **(E)** side views. Superimpositions of the transmembrane domains (S1-S6) were created using Fr-TM-Align.

**Fig. S5.**
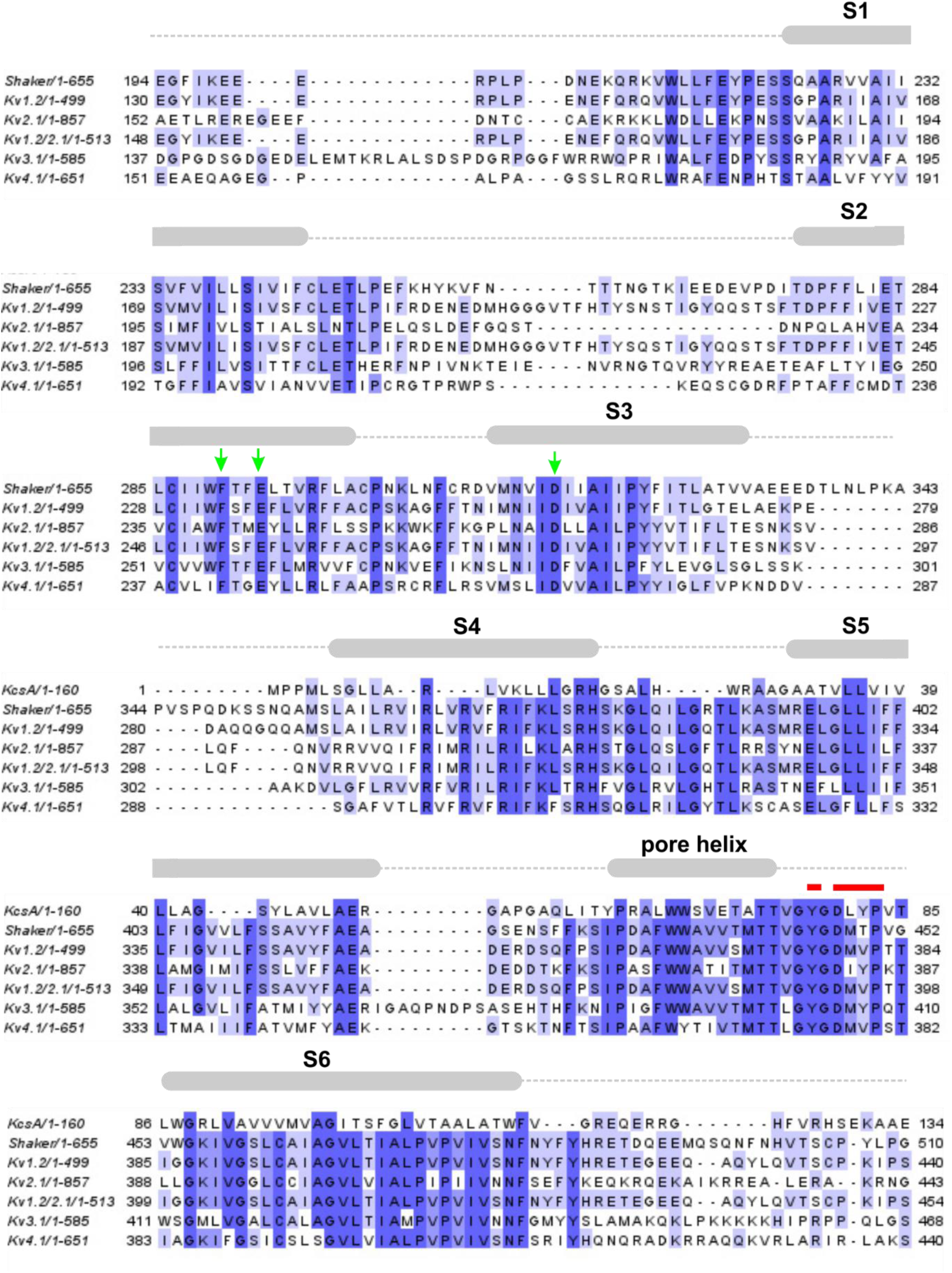
Sequence alignment of Kv channels. Sequence alignment of Shaker (*Drosophila melanogaster* P08510), KcsA (*Streptomyces lividans* P0A334), Kv1.2 (*Rattus norvegicus* P63142), Kv2.1 (*Rattus norvegicus* P15387), Kv1.2/2.1 chimera(*Rattus norvegicus* PDB: 6EBK), Kv3.1 (*Rattus norvegicus* P25122) and Kv4.1 (*Mus Musculus* Q03719). Blue regions indicate similarity and dark blue region indicate identity. Cartoons represent secondary structure features. Green arrows highlight important residues in the chargetransfer center and red bar indicates residues that change conformation most dramatically between Shaker-IR and W434F.

**Fig. S6.**
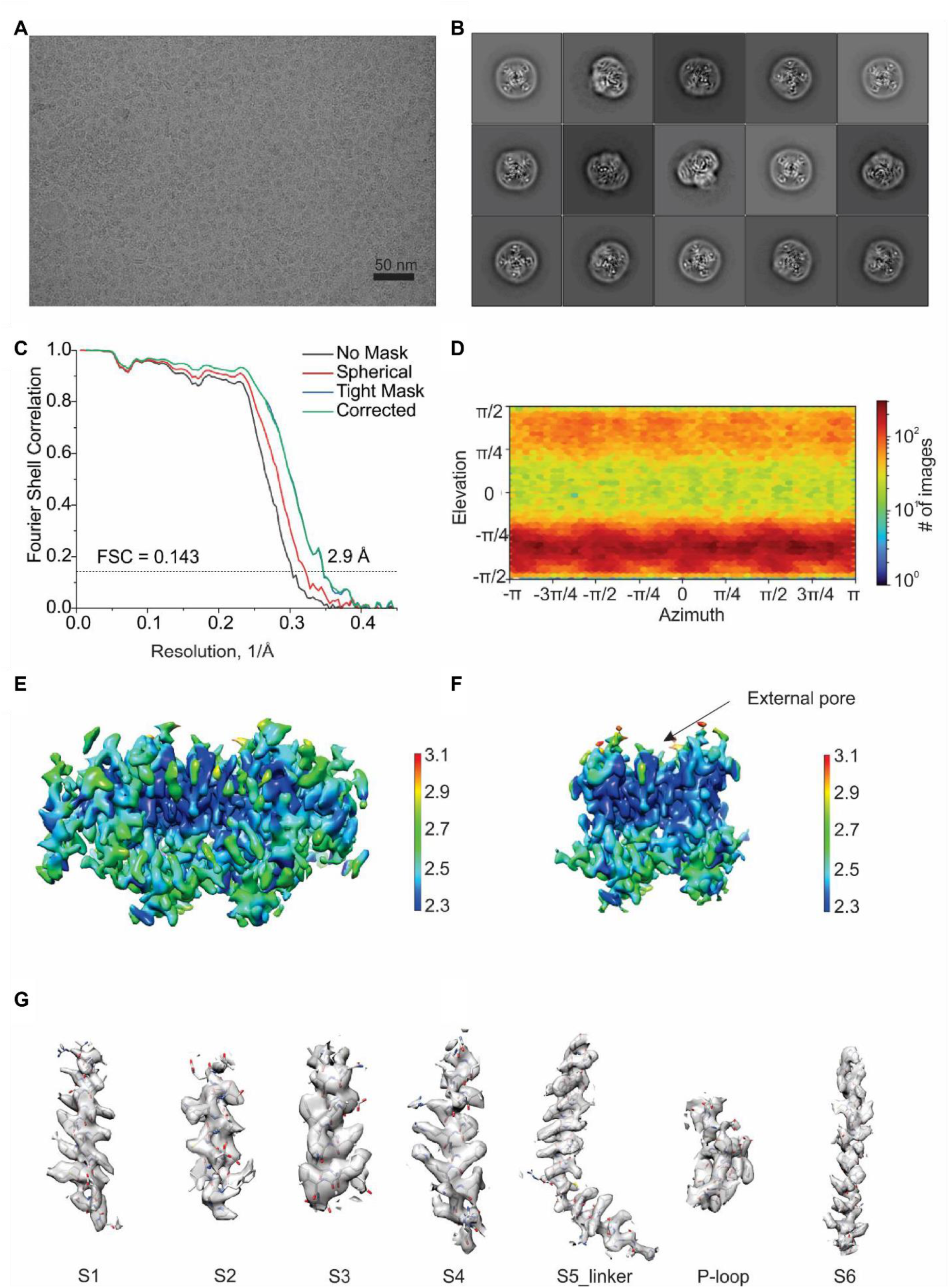
Cryo-EM imaging of Shaker-IR W434F. **(A)** Cryo-EM micrograph of Shaker-IR W434F. **(B)** 2D class averages of the particles in different orientations. **(C)** Fourier Shell Correlation (FSC) curves. **(D)** Direction distribution plots of the 3D reconstruction. **(E)** Local resolution map for the entire TM region. **(F)** Local resolution map for the S5-S6 pore domain, highlighting the dark blue region within the outer pore that shows the best resolution in the overall structure. **(G)** Regional cryo-EM density for Shaker-IR W434F.

**Fig. S7.**
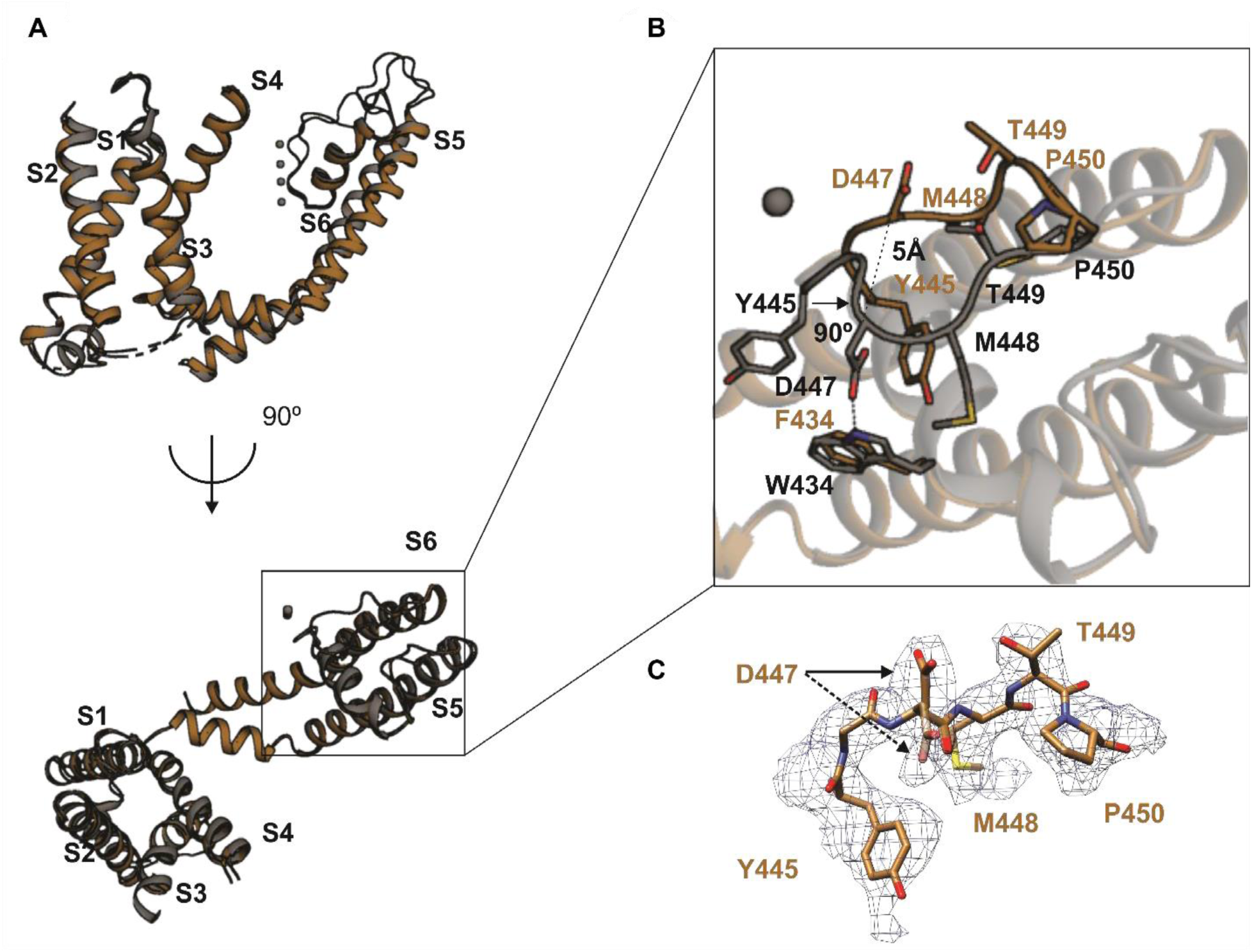
Structural alignment of Shaker-IR with the W434F mutant. **(A)** Superimposition of the Shaker-IR (gray) and W434F (brown) viewed from the side (top image)or from the external side of the membrane (bottom). **(B)** Close-up view of the conformational change within the outer pore domain viewed from the external side of the membrane. **(C)** Cryo-EM density of P-loop for the W434F mutant. Density for two distinct rotamers of D447 are indicated by arrows.

**Fig. S8.**
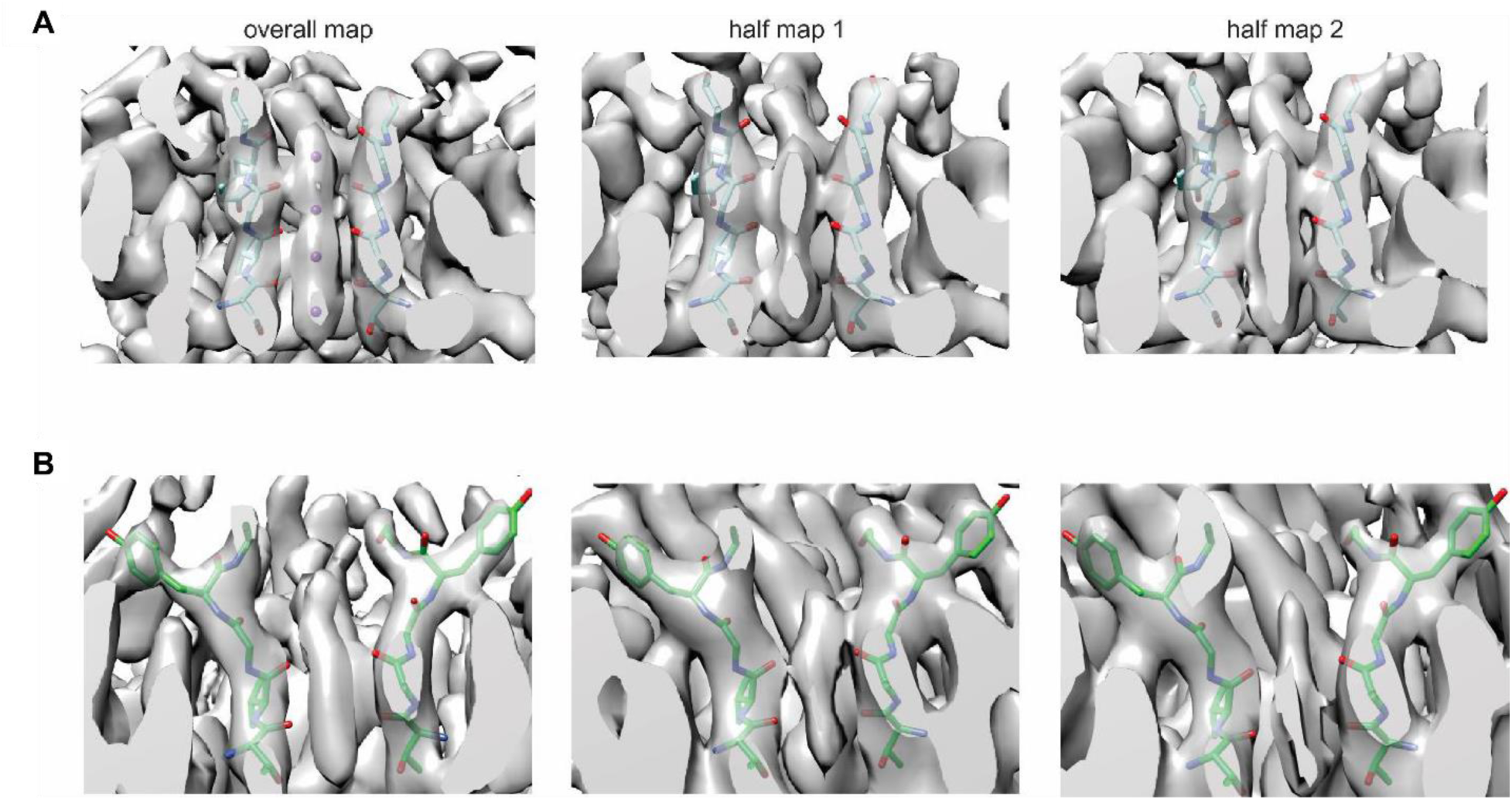
Ion densities in the selectivity filter of the wild-type Shaker-IR channeland W434F mutant. **(A)** Ion density of Shaker-IR using C1 symmetry reconstruction shown in the overall map and half maps (lowpass at 3.5 Å). 4 ions (purple) are fit into S1-S4 sites (left panel), with residues in the filter shown in stick. **(B)** Ion density of Shaker-IR W434F using C1 symmetry reconstruction shownin the overall map and half maps (lowpass at 3.5 Å). Cryo-EM density between S1 and S2 sites inW434F are much weaker compared the Shaker-IR.

**Fig. S9.**
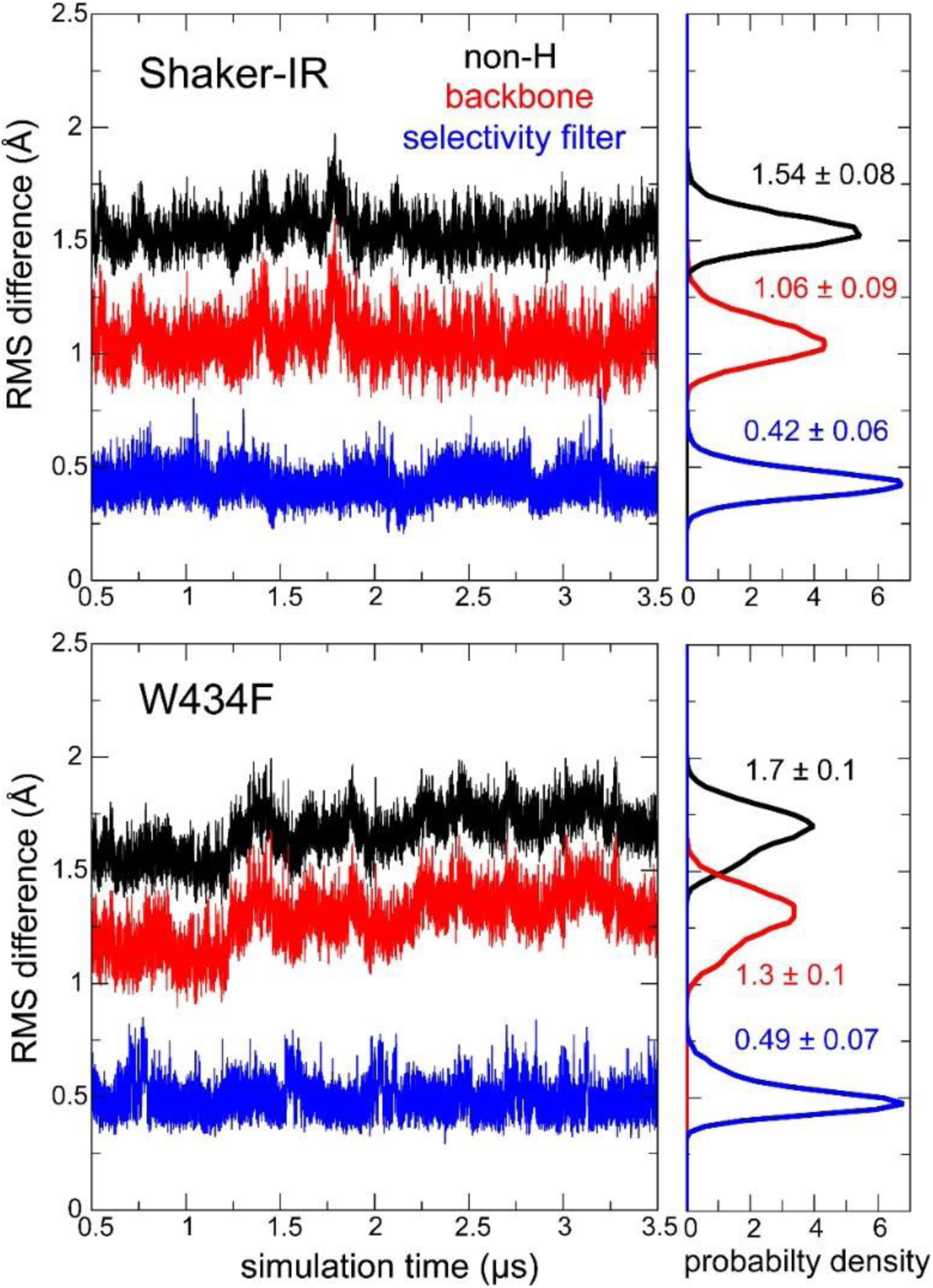
MD simulations of Shaker-IR and the W434F mutant. For the portion of the calculated MD trajectories wherein a transmembrane voltage is applied (300 mV for Shaker-IR and 300/450 mV for W434F), the figures on the left side quantify the root-mean-square (RMS) difference between each of the snapshots (in intervals of 120 ps) and the corresponding cryo-EM structure. RMS differences are quantified for all non-hydrogen atoms in the protein structures (black), only for the backbone atoms (red), and only for the backbone atoms of residues 442 to 445. On the right, the figure shows normalized probability histograms of the time-series shown on the left; mean values and standard-deviations are provided in each case.

**Fig. S10.**
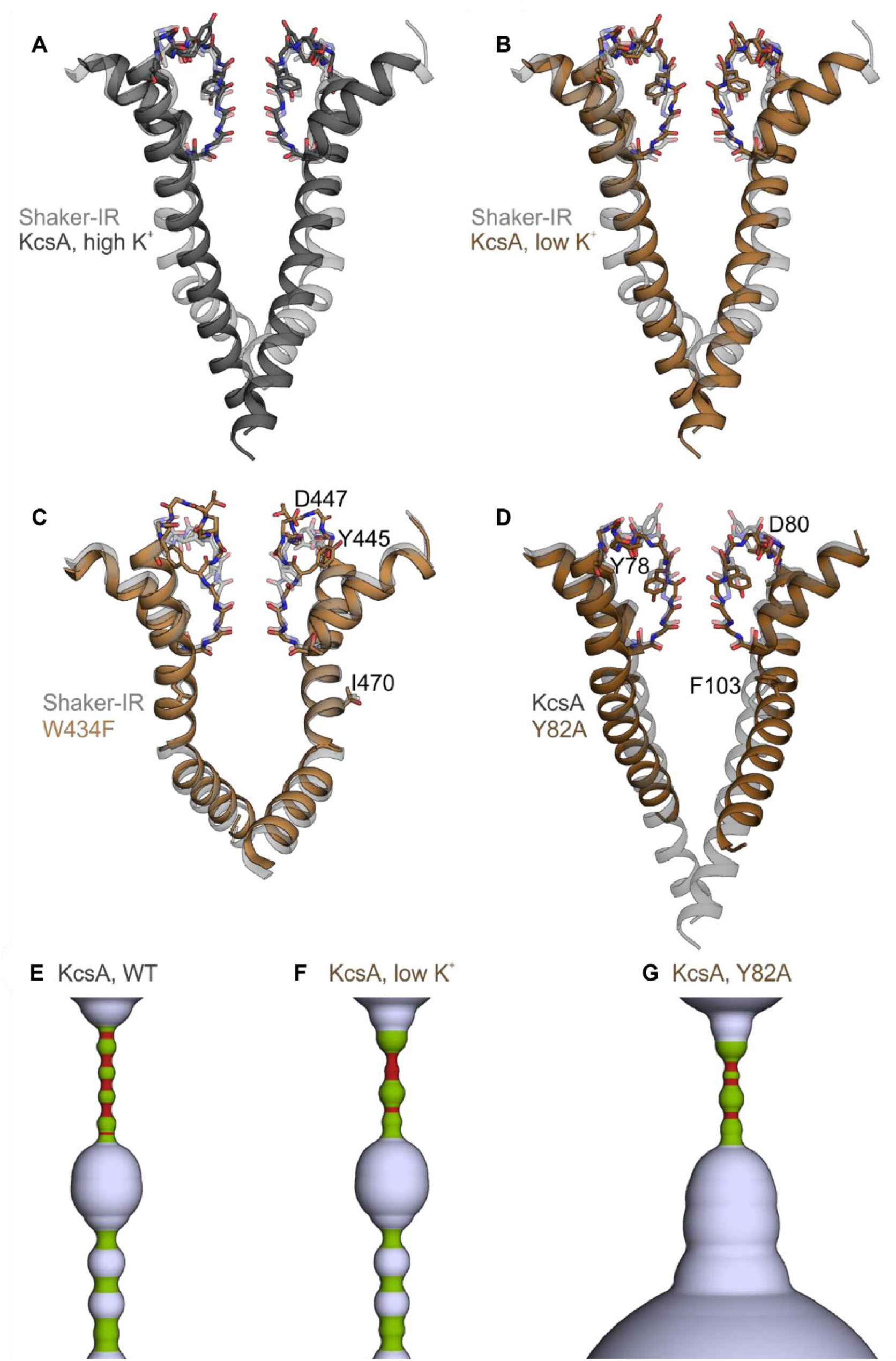
Comparison of Shaker Kv channel structures to KcsA. **(A)** Superimposition of S6 and pore helix (two opposing subunits) of Shaker-IR (7SIP) and KcsA inthe presence of high K^+^ (200 mM K^+^, 1K4C). **(B)** Superimposition of S6 and pore helix of Shaker-IR (7SIP) and KcsA in the presence of low K^+^ (3 mM K^+^, 1K4D). **(C)** Superimposition of S6 and pore helix of Shaker-IR (7SIP) and W434F mutant (7SJ1). **(D)** Superimposition of S6 and pore helix of KcsA in the presence of high K^+^ (1K4C) and KcsA fast-inactivating Y28A mutant in the inactivated/open state (5VKE). Superimpositions of the pore domain (S5-S6) were created using Fr-TM-Align. HOLE diagrams of **(E)** KcsA in the presence of high K^+^ (1K4C), **(F)** KcsA in the presence of low K^+^ (1K4D), and **(G)** KcsA fast-inactivating Y28K mutant (5VKE). Radii ≤ 1Å are shown in red, radii ≤ 2Å and > 1Å are shown in green, and radii larger than 2Å are shown in light blue.

**Table S1.** Cryo-EM data collection, refinement and validation statistics.

|  | Shaker-IR<br>(EMDB-25147)<br>(PDB 7SIP) | Shaker-W434F<br>(EMDB-25152)<br>(PDB 7SJ1) |
| --- | --- | --- |
| **Data collection and processing** |  |  |
| Magnification | 105,000 | 105,000 |
| Voltage (kV) | 300 | 300 |
| Detector | Gatan K3 | Gatan K3 |
| Electron exposure (e-/Å<sup>2</sup>) | 52 | 52 |
| Defocus range (μm) | -0.5 to -1.5 | -0.5 to -1.5 |
| Pixel size (Å) | 0.43 | 0.43 |
| Symmetry imposed | C4 | C4 |
| Initial particle images (no.) | 4,934,881 | 7,446,104 |
| Final particle images (no.) | 229,379 | 212,083 |
| Map resolution (Å) | 3.0 | 2.9 |
| FSC threshold | 0.143 | 0.143 |
| **Refinement** |  |  |
| Initial model used (PDB code) | 6EBM | 6EBM |
| Model resolution (Å) | 3.1 | 3.2 |
| FSC threshold | 0.5 | 0.5 |
| Map sharpening *B* factor (Å<sup>2</sup>) | -129 | -102 |
| Model composition |  |  |
| Non-hydrogen atoms | 7560 | 6495 |
| Protein residues | 892 | 784 |
| Ligands | 36 | 35 |
| *B* factors (Å<sup>2</sup>) |  |  |
| Protein | 65.23 | 35.3 |
| Ligand | 60.7 | 28.7 |
| R.m.s. deviations |  |  |
| Bond lengths (Å) | 0.003 | 0.003 |
| Bond angles (°) | 0.508 | 0.531 |
| Validation |  |  |
| MolProbity score | 1.14 | 1.43 |
| Clashscore | 3.49 | 7.17 |
| Poor rotamers (%) | 0 | 0 |
| Ramachandran plot |  |  |
| Favored (%) | 98.72 | 98.01 |
| Allowed (%) | 1.28 | 1.99 |
| Disallowed (%) | 0 | 0 |

**Table S2.** Multistep protocol to equilibrate the molecular systems constructed in this simulation study.

| Step <sup>a</sup> | 1 | 2 | 3 | 4 | 5 | 6 | 7 | 8 | 9 |
| --- | --- | --- | --- | --- | --- | --- | --- | --- | --- |
| Positional restraints <sup>b</sup><br>(kcal/mol/Å<sup>2</sup>) | 60 | 60 | 15 | 4 | - | - | - | - | - |
| Distance restraints <sup>c</sup><br>(kcal/mol/Å<sup>2</sup>) | - | - | - | - | 4 | 1 | 1 | - | - |
| Dihedral $\Phi$, $\Psi$ and $\chi_1$ restraints<br>(kcal/mol/deg<sup>2</sup>) | - | - | - | - | 16 | 4 | 4 | 4 | 1 |
| Distance-to-center restraints <sup>d</sup><br>(kcal/mol/Å<sup>2</sup>) | - | - | - | - | 60 | 15 | - | - | - |
| Center-of-mass restraints <sup>e</sup><br>(kcal/mol/Å<sup>2</sup>) | 1 | 1 | 1 | 1 | 1 | 1 | 1 | 1 | 1 |
| Length of simulation (ns) | 1 | 1 | 1 | 1 | 50 | 25 | 25 | 25 | 25 |

<sup>a</sup> This protocol was carried out with NAMD 2.13 at 0 mV and preceded the trajectories calculated on Anton2 discussed in the manuscript. All restraints used during the equilibration are harmonic potentials. The table specifies the force constant used for each kind at each step in the protocol.
<sup>b</sup> Positional restraints on all non-hydrogen atoms in the protein and on K<sup>+</sup> and water oxygens in the selectivity filter.
<sup>c</sup> Distance restraints on selected hydrogen-bonds and hydrophobic interactions behind the selectivity filter.
<sup>d</sup> K<sup>+</sup> and water molecules were restrained to remain near the geometric center of their designated binding sites in selectivity filter; each center was defined by eight backbone/sidechain oxygens.
<sup>e</sup> The protein center-of-mass was restrained to remain near the center of the simulation box.

**Movie S1.**

**Morph between the structures of Shaker-IR and the W434F mutant**

**Movie S2.**

**MD simulation of K^+^ dynamics and permeation through Shaker-IR.** The movie depicts snapshots of K^+^ (magenta, red, purple, yellow, orange spheres) near and withinthe selectivity filter of Shaker-IR (gray cartoon), for a 1-µs fragment of the trajectory (starting at 1.5 µs), under 300 mV. Water molecules with 3.8 Å of these K^+^ ions are shown (cyan spheres). The backbone atoms of residues 443-445 and the backbone and sidechain atoms of residue 442are highlighted (excluding hydrogens).

**Movie S3.**

**MD simulation of K^+^ dynamics and permeation through Shaker-W434F.** The movie depicts snapshots of K^+^ (magenta, red, purple, yellow, orange spheres) near and withinthe selectivity filter of Shaker-W434F (gray cartoon), for a 1-µs fragment of the trajectory (starting at 2.0 µs), under 300 mV for the first 500 ns and under 450 mV for the last 500 ns. Water molecules within 3.8 Å of these K^+^ ions are shown (cyan spheres). The backbone atoms of residues 443-445 and the backbone and sidechain atoms of residue 442 are highlighted (excluding hydrogens).

## Notes

### Competing Interest Statement

The authors have declared no competing interest.

